# Cryo-EM Structures of Four Polymorphic TDP-43 Amyloid Cores

**DOI:** 10.1101/601120

**Authors:** Qin Cao, David R. Boyer, Michael R. Sawaya, Peng Ge, David S. Eisenberg

## Abstract

TDP-43 is an essential DNA/RNA processing protein that undergoes both functional and pathogenic aggregation. Functional TDP-43 aggregates are reversible, forming transient species such as nuclear bodies, stress granules, and myo-granules^1–3^. In contrast pathogenic TDP-43 aggregates are irreversible, forming stable intracellular amyloid-like inclusions^4,5^. These inclusions are the primary pathology of amyotrophic lateral sclerosis (ALS) and frontotemporal lobar degeneration with TDP-43 inclusions (FTLD-TDP)^6^. Disease-associated, hereditary mutations in TDP-43 are known to accelerate the deposition of irreversible aggregates in the cytoplasm^7^. Reversible TDP-43 aggregation has been shown to precede the formation of irreversible amyloid fibrils similar to the behavior of proteins hnRNPA1 and FUS^8–10^. Still unknown, however, are the structural features of TDP-43 fibrils that confer both reversibility and irreversibility and how hereditary mutations can impose irreversible aggregation. Here, we determined the structures of amyloid fibrils formed by two segments previously reported to be the pathogenic cores of TDP-43 aggregation^7,11,12^; these are termed SegA (residues 311-360) and SegB A315E (residues 286-331 containing the ALS hereditary mutation A315E). SegA forms three polymorphs, all with dagger-shaped folds. SegB forms R-shaped folds. All four polymorphs have folds confined to two dimensions, and are stabilized by hydrophobic cores and peripheral hydrogen bonds. Energetic analysis suggests that the dagger-shaped polymorphs are examples of the irreversible fibril structures of TDP-43, whereas the SegB polymorph may participate in both reversible and irreversible fibril structure. Our structure suggests how the A315E mutation may convert this polymorph to the irreversible type and lead to mutation-enhanced pathology.

## Main text

Amyloid-forming proteins seem to violate the central tenant of protein science—that amino acid sequence determines structure and function^13^. In contrast to globular and membrane proteins each of which folds into a single functional structure, a given amyloid-forming sequence can fold into several distinctly different polymorphic structures^14,15^. Here we find that TDP-43, known to form both reversible functional aggregates and irreversible pathogenic aggregates, exhibits polymorphic behavior. This complex protein thus offers an intriguing opportunity for structural exploration.

Irreversible, hyperphosphorylated aggregates of C-terminal segments of TDP-43 are found in the autopsied brains of ALS and FTLD-TDP patients^6,16,17^. These aggregates have also been found in Alzheimer’s, Parkinson’s, CTE, Huntington’s disease and inclusion body myopathies (IBMs), among others^18–21^. Because TDP-43 functions in several essential steps of RNA metabolism^22,23^, it is widely considered that TDP-43 aggregation is toxic through a loss-of-function mechanism^24–26^. Structural studies of amyloid fibrils of β-amyloid^27,28^, tau^29,30^, α-synuclein^14^, and β2-microglobulin^31^ have revealed polymorphs and insights into pathogenesis. Here, we use cryo-EM to determine the overall folds of TDP-43 amyloid cores, expanding structural information beyond the local interactions previously revealed by crystallography^32,33^.

We obtained TDP-43 fibrils by incubating SUMO-tagged TDP-43 segments in the test tube after the cleavage of SUMO tag (Supplementary Figure 1a). We first aimed to produce fibrils formed by full-length TDP-43, a pathogenic C-terminal fragment (CTF, 208-414) – truncation product that is enriched in disease brain^34^, or by the Low Complexity Domain (LCD, 274-414) which is considered to be necessary for TDP-43 aggregation^32,35,36^. However, despite our efforts at optimization, we could observe only highly clumped fibril-like structures and/or disordered aggregates that are not suitable for cryoEM structure determination (See Supplementary Figure 1b top panel). We suspect that this may be due to the ability of longer constructs of TDP-43 to participate in multi-valent interactions, possibly through LARKS^32,37^ or other adhesive segments outside the LCD^33^. These multi-valent interactions have been shown to assemble networks of protein chains and could conceivably explain why longer segments of TDP-43 form amorphous aggregates or fibril clumps not amenable to cryoEM structure determination. This observation is in line with TDP-43’s role in phase separation and stress granule formation, which requires the presence of multi-valent interactions^37^.

To overcome the hurdle of the disordered assembly of longer segments of TDP-43, we implemented a “divide and conquer” approach whereby we selected known aggregation cores for structure determination. We chose SegA (residues 311-360) and SegB (residues 286-331), guided by the following evidence. SegA was previously identified as an aggregation core of TDP-43 due to the observation that its deletion decreases TDP-43 aggregation in vitro and in cells, whereas addition of SegA to the aggregation-resistant *C. elegans* TDP-43 homolog induces aggregation^11^. Fibrils of SegB are toxic to primary neurons, and an ALS hereditary mutation A315T together with phosphorylation of the threonine, which is speculated to occur in hyper-phosphorylated aggregates of TDP-43 in disease, increases SegB’s cytotoxicity^7^. With this in mind, we selected SegB A315E – another hereditary mutation with similar effects as A315T^7,38–40^ and a mimic of A315T with phosphorylation – in order to visualize the structure of a second possible TDP-43 aggregation core and to gain insight into the molecular mechanism of mutation-enhanced TDP-43 pathology. The importance of SegA and SegB in full-length TDP-43 aggregation is also supported by other studies which found that amyloid fibrils containing either a core region (residues 314-353) of SegA or region (residues 274-313) similar to SegB can template aggregation of full-length TDP-43 in SH-SY5Y human neuronal cells^41^. Likewise, in the same cell line, deletion of these two regions (residues 314-353 or 274-313) from full-length TDP-43 inhibits aggregation. As we expected, fibrils formed by SegA and SegB A315E were much more homogenous and less bundled than longer segments of TDP-43, including SegAB (286-360) that contains both aggregation cores SegA and SegB (Supplementary Figure 1b). This observation supports the idea that eliminating competing multi-valent interactions helps to produce homogenous fibrils of isolated amyloid cores. Using these homogenous preparations, we determined three polymorphic fibril structures of SegA (termed SegA-sym, SegA-asym, and SegA-slow) and one of SegB A315E (Figure 1, Supplementary Figure 2-4 and Supplementary Table 1, see Methods for technical details).

**Figure 1.**
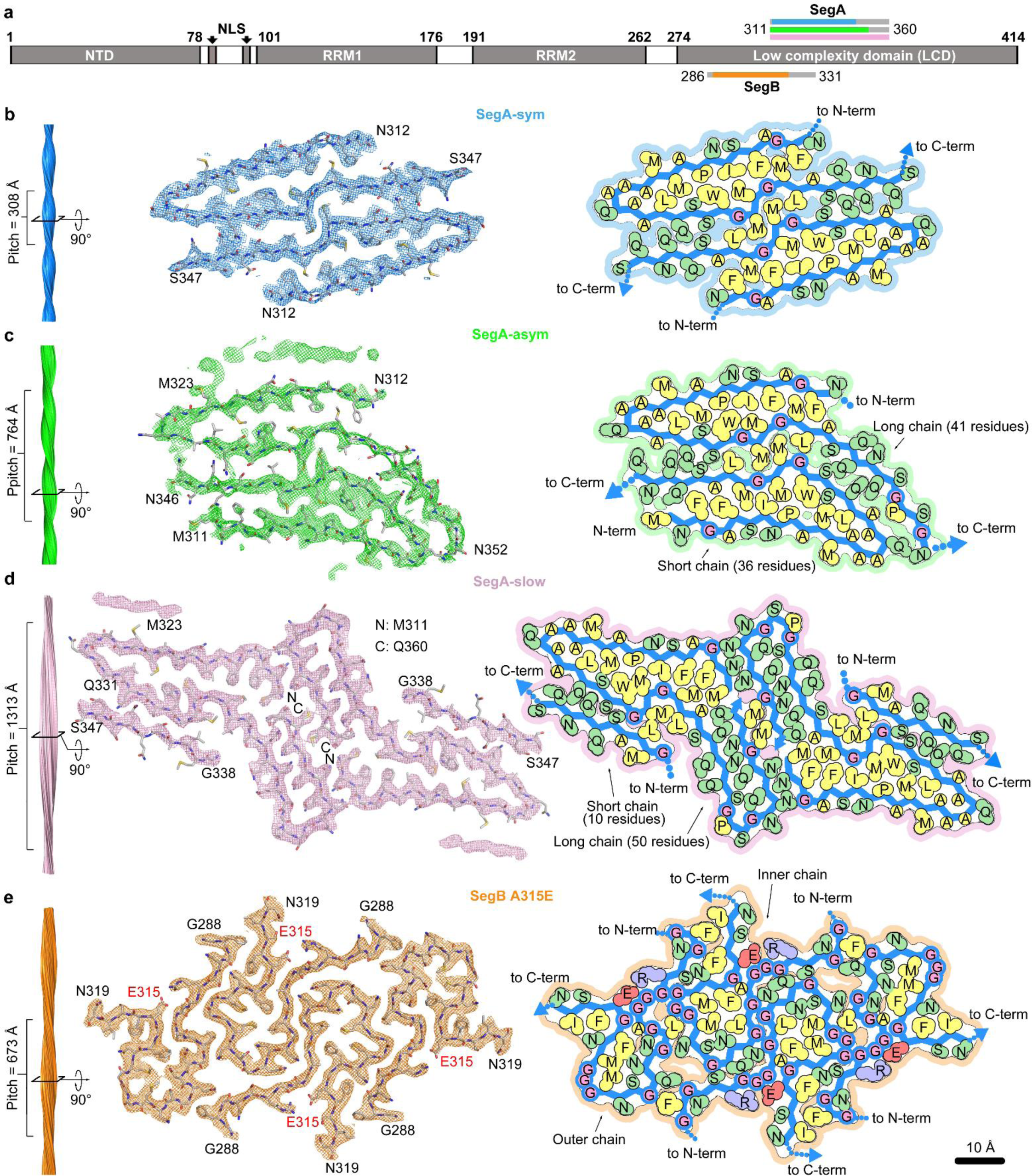
Cryo-EM Structure of TDP-43 polymorphic fibrils. **a**, Schematic of full-length TDP-43. SegA (residues 311-360) and SegB (residues 286-331) identified for structural determination are shown as gray bars, respectively above and below the low-complexity domain (LCD). The color bars show the range of residues visualized in the structure of each polymorph. Panels **b**-**e**: (left) fibril reconstructions showing left-handed twist and pitch; (middle) density and atomic model of one cross-sectional layer of each fibril; (right) schematic model showing protein chain (blue) and residues (hydrophobic in yellow, polar in green, glycine in pink, glutamate in red, and arginine in blue).

All four fibrils are formed from gradually twisting β-sheets that run the entire length of the fibrils (Figure 1b-e). A thin slab or “layer” perpendicular to each fibril axis (Figure 1, middle) shows that individual SegA and SegB chains are each confined within an essentially two-dimensional layer, in contrast to globular and membrane proteins whose folds occupy three-dimensions. Identical layers stack on one another creating β-sheets that are parallel and in-register. Each layer contains two or more protein chains, giving rise to a corresponding number of protofilaments in the fibril.

SegA polymorphs all share a dagger-shaped fold comprised of residues 312-346 and a sharp 160° kink at Gln327, which forms the dagger tip (Figure 2a-d, Supplementary Figure 5a-c, detailed comparison see Supplementary Note 1&2). The three SegA polymorphs mainly differ in the number of protofilaments and symmetry. SegA-sym fibrils contain two protofilaments related by a pseudo-2_1_ axis, whereas SegA-asym contains two protofilaments with somewhat different conformations. SegA-slow fibrils contain four protofilaments; two protofilaments are related by a central 2-fold axis and contain 50 ordered residues. These are flanked by two other protofilaments containing only 10 ordered residues. We note that SegA-sym and SegA-asym structures are compatible with full-length TDP-43 due to their free N-and C-termini, whereas the SegA-slow structure is an artifact of the truncation since the N- and C-termini are sequestered in the center of the fibrils (Figure 1d and Supplementary Figure 5b). Therefore, in the following analysis, we mostly focus on SegA-sym and SegA-asym. However, we note that SegA-slow provides valuable information, including validation of the dagger-shaped fold owing to its higher resolution and direct atomic evidence of secondary nucleation (Supplementary Note 3).

**Figure 2.**
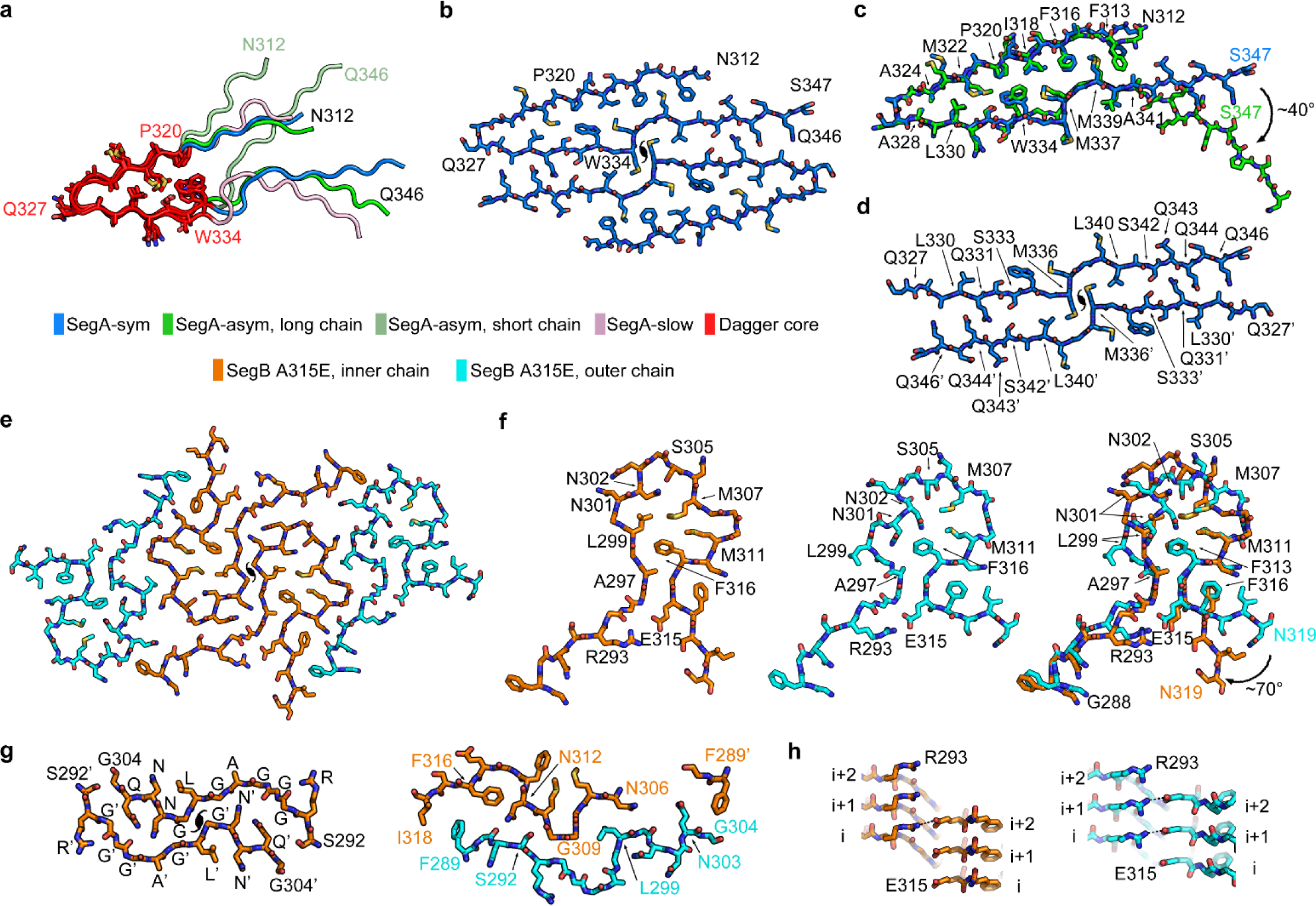
Structure of the dagger-shaped fold and the R-shaped fold. **a**, Superposition of the dagger-shaped folds (residue 312-346) in three SegA polymorphic fibrils. The core region of the dagger-shaped folds (residue 320-334) is colored red with side chains shown, and the rest of the dagger-shaped fold is colored according to the key. **b**, Atomic model of SegA-sym. **c**, Pairwise superposition of the dagger-shaped fold of SegA-sym vs. SegA-asym long chain. **d**, Continuous dimer interface of SegA-sym. **e**, Atomic model of SegB A315E. Each of the four chains is R-shaped, with a pseudo-2_1_ axis relating the two pairs. The inner two chains are identical as are the two outer chains. **f**, Structures of the inner (left) and outer (middle) chains and their superposition (right). Notice the proximity of R293 to the pathogenic variant E315 which replaces A315 in the reference sequence. **g**, View parallel to the fibril axis showing the symmetric interface between the two inner chains (left) and the asymmetric interface between the inner and outer chains (right). **h**, View perpendicular to the fibril showing the inter-layer interaction of Arg293 with Glu315 on the inner chain (left) and outer chain (right). (Detailed alignment parameters and RMSD values listed in Supplementary Table 2).

SegB A315E forms fibrils of a single morphology, in contrast to polymorphic SegA. These fibrils are characterized by an R-shaped fold spanning residues 288-319 (Figure 2e&f and Supplementary Note 4). Overall, the R-shaped fold is more highly kinked than the dagger-shaped fold (Figure 1b-e), probably due to the abundance of glycine residues. Each fibril is wound from four protofilaments (Figure 2e). The two inner protofilaments are related through a tight, pseudo-2_1_ symmetric interface (Figure 2e&g, Supplementary Note 5). These protofilaments are flanked by two outer protofilaments through an asymmetric interface involving outer residues 289-304 and inner residues 289 and 306-318 (Figure 2e&g, Supplementary Note 5). Small conformational differences between inner and outer R-folds (Figure 2f) recall the positional polymorphism observed in a different, shorter TDP-43 segment^33^.

In all four R-folds we observe a salt-bridge between Arg293 and Glu315 enabled by the pathogenic A315E mutation. The Arg293-Glu315 salt-bridge is not formed by the residues from the same layer along the fibril axis. Rather, each Arg293 interacts with the Glu315 from one or two layers above (Figure 2h), which may affect the kinetics of fibril growth and nucleation. This salt-bridge suggests a mechanistic explanation for the inclination of A315E toward TDP-43 pathology. Model building suggests that wild-type SegB can form the same R-shaped fold as A315E, with Ala315 participating in a hydrophobic interaction with Ala297 and Phe313. This hypothesis is supported by the similar stability and morphology of wild-type SegB fibrils in our themostability assays (Supplementary Figure 1c) compared to SegB A315E fibrils and is also supported by the cross-seeding ability of SegB and SegB A315E (Supplementary Figure 1e).

It is noteworthy that although TDP-43 can form either the dagger-shaped polymorph of SegA or the R-shaped polymorph of SegB, it is unlikely that a given molecule of TDP-43 can form both simultaneously. This exclusivity is indicated by a superposition of the two folds in the overlapping segment Asn312-Asn319 (Supplementary Figure 5h). The superposition reveals incompatible steric hindrance between the main chains of SegA and SegB would occur if both folds were formed by a single TDP-43 molecule that contains both SegA and SegB sequences.

The extensive hydrophobic interactions and hydrogen bonds observed in both the dagger-and R-shaped folds suggest that both SegA and SegB A315E fibrils are irreversible. To examine this hypothesis, we calculated modified atomic solvation energies (see Methods) to quantify the stabilities of the fibrils. The calculated stabilities of both the dagger-and R-shaped fold fibrils (represented by energy per layer and per residue) are comparable to other pathogenic amyloid fibrils, such as the Alzheimer’s and Pick’s disease fibrils (Figure 3). In contrast the stability of FUS fibrils, thought to be reversible^42^, is distinctly lower. We performed thermostability assays to validate our energetic calculations. When heated to 75 °C for 30 minutes, fibrils formed by SegA, SegB, and SegB A315E are all stable, whereas fibrils formed by mCherry-FUS-LCD (composed of residues 1-214 – identical to the sequence determined in the FUS ssNMR structure except with a His-tag replacing mCherry^42^) disappeared after heating to 60 °C (Supplementary Figure 1c). These results are consistent with our energetic calculations and support the idea that the dagger-and R-shaped fold fibrils are irreversible.

**Figure 3.**
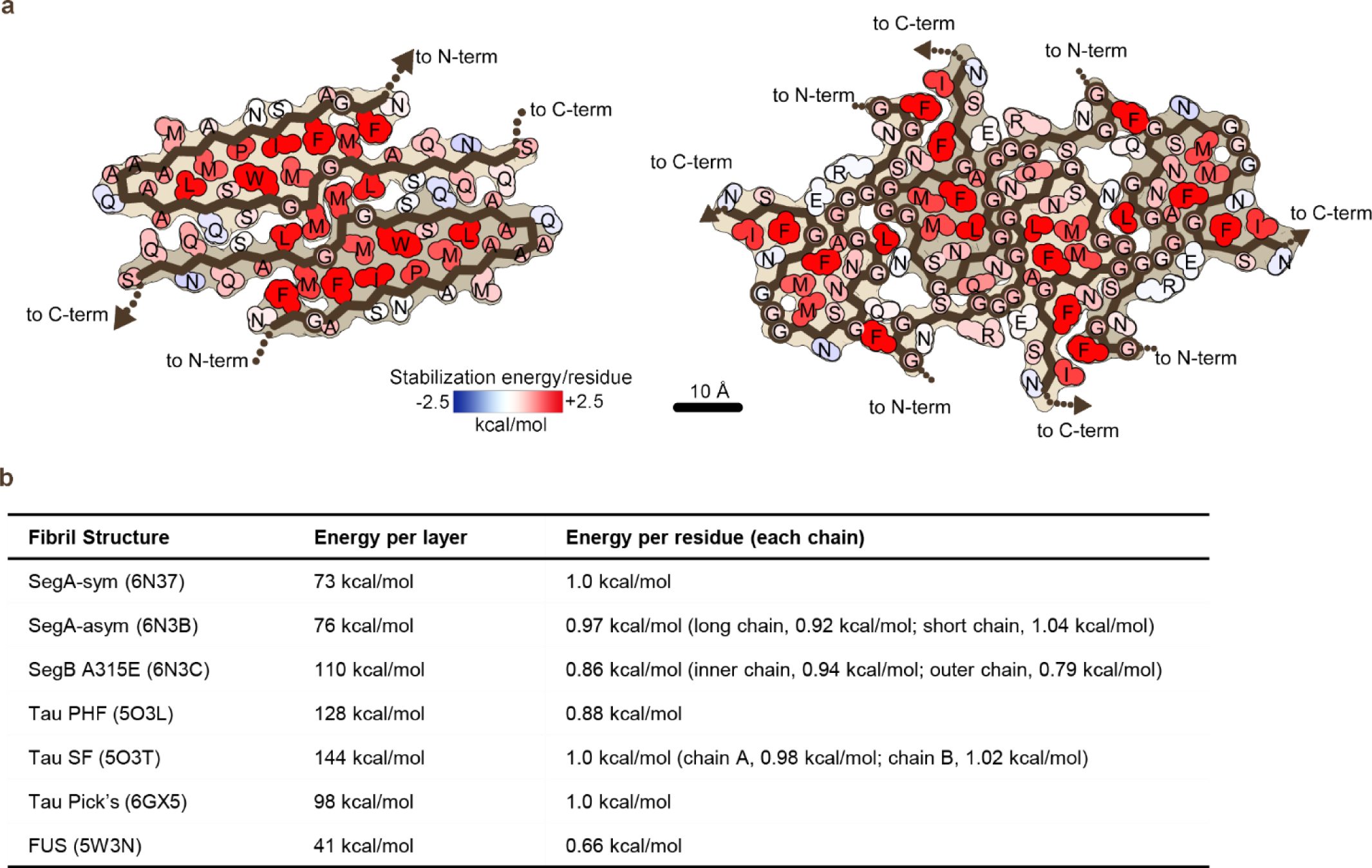
Calculated solvation energy maps for TDP-43 fibrils based on atomic solvation parameters. **a**, Solvation energy maps of SegA-sym (left) and SegB A315E (right). Residues are colored from unfavorable (blue, −2.5 kcal/mol) to favorable stabilization energy (red, 2.5 kcal/mol). Notice that both the dagger-shaped fold (SegA-sym) and the R-shaped fold (SegB A315E) show stable cores. **b**, Comparative solvation energy calculations. Stabilization energies of seven fibrils are listed as energy per layer as shown here (representing overall stability of each fibril), per chain and per residue. Notice that the overall stability of the dagger-shaped fold (SegA-sym and SegA-asym) and the R-shaped fold (SegB A315E) are comparable with three brain extracted tau fibrils, and more stable than FUS fibrils that are considered to representative of reversible aggregates.

Several lines of evidence suggest that the dagger and R-shaped folds can be adopted by longer TDP-43 segments such as the pathogenic TDP-CTF fragment. (1) Structural conservation: The core region (residue 320-334) of the dagger-shaped fold is structurally conserved in all four dagger-shaped folds despite differences in local environment (Supplementary Note 1). (2) Model building. The outward facing disposition of the termini in these folds longer TDP-43 structures to be modeled from these cores without steric interference. (3) Cross seeding. Fibrils formed by TDP-CTF can seed SegA and SegB A315E monomer (Supplementary Figure 1f), and SegA and SegB A315E fibrils can seed TDP-LCD monomer (Figure 4b) (due to its rapid aggregation, TDP-CTF monomer could not be used to measure seeding and we instead used TDP-LCD monomer as the LCD is required for TDP-43 aggregation^32,35,36^). (4) Mutagenesis experiments with TDP-CTF reported here and previously^32^. We found that five individual point mutations to tryptophan delay aggregation of TDP-CTF; these sterically conflict with the tightly packed dagger-shaped (A324W, L330W, Q331W, M337W) and R-shaped (Q303W) folds delay aggregation of TDP-CTF. Similarly, the mutations A324E and M337E that are disruptive to the dagger-shaped fold are found to inhibit aggregation of full-length TDP-43 in cells^11^. As negative controls, we found that A326W and G304W – located in a solvent exposed environment of the dagger-shaped fold and a loose cavity of the R-shaped fold, respectively – can tolerate the bulky tryptophan and did not delay aggregation of TDP-CTF (Supplementary Figure 1d, Supplementary Figure 6a&b, Supplementary Figure 7 and Supplementary Note 5)^32^. Furthermore, a key stabilizing feature of SegB A315E, the Arg293-Glu315 salt bridge, is similarly important for TDP-CTF aggregation. We designed two double mutations, R293E/A315E and R293E/A315R. Based on our structure of SegB A315E, R293E/A315E would disrupt the R-shaped fold by electrostatic repulsion whereas R293E/A315R would favor the R-shaped fold by restoring the salt-bridge. In accordance with our model, R293E/A315E reduces TDP-CTF aggregation whereas R293E/A315R enhances TDP-CTF aggregation (Supplementary Figure 1d). These results suggest that both the dagger-and the R-shaped folds are accessible to TDP-CTF molecules, so that disrupting either one blocks one pathway to fibril formation and hence delays aggregation of TDP-CTF. The observation that none of these mutations targeting the dagger-or R-shaped fold fully eliminates aggregation supports our hypothesis that TDP-CTF can form fibrils using one of multiple aggregation cores, including the dagger-or R-shaped fold, or possibly others not discovered here. In short, conservation of the dagger fold, modeling, seeding experiments, and mutational analysis all suggest that TDP-CTF can form the folds we observe.

**Figure 4.**
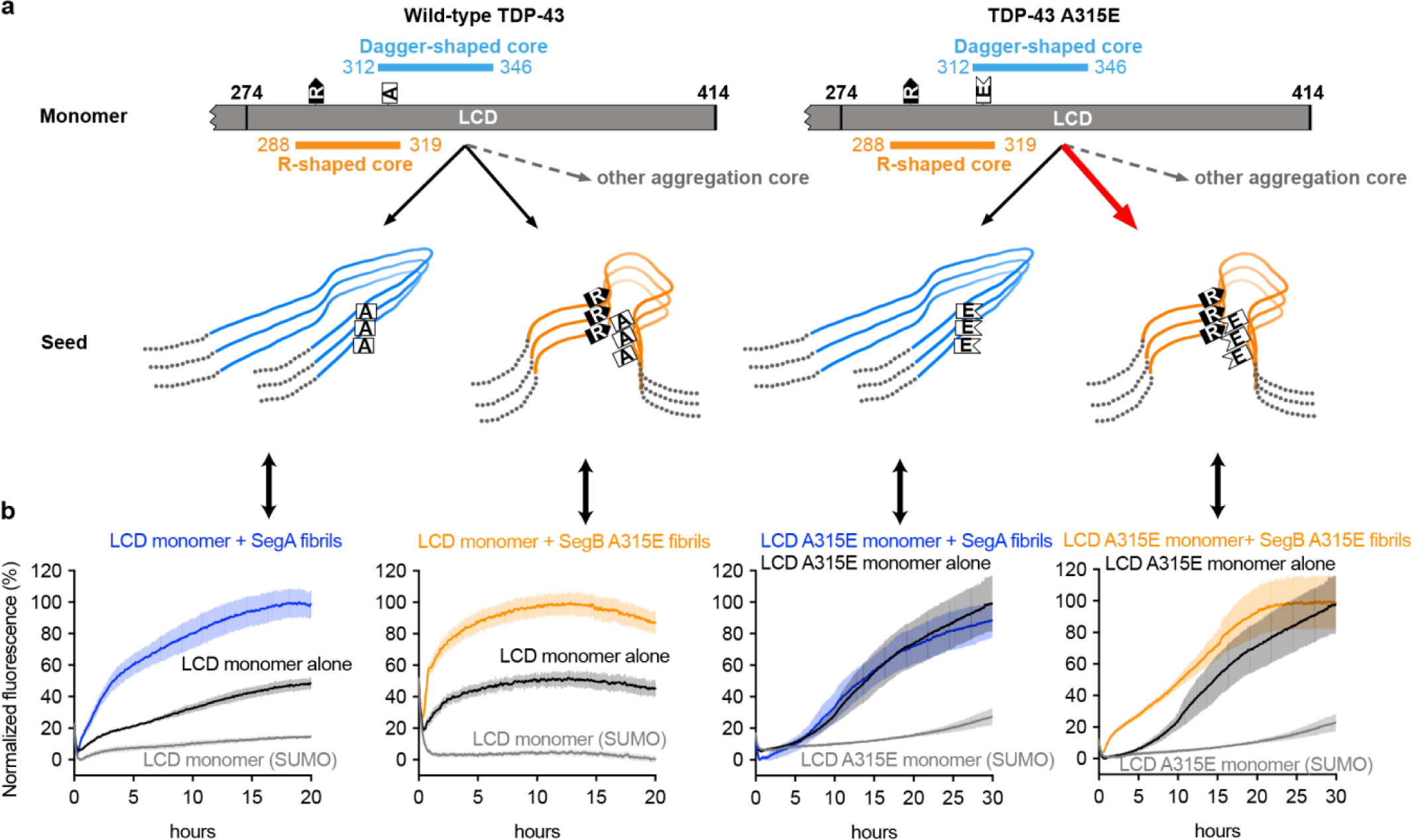
Speculative model of TDP-43 amyloid aggregation based on fibril structures reported here. **a**, (Left) Monomeric TDP-43 can form amyloid fibrils with either dagger-shaped or R-shaped cores (or perhaps other so far unidentified cores). The ALS hereditary A315E mutation accelerates the R-shaped core through electrostatic interaction of Glu315 with Arg293. **b**, Experiments seeding TDP-43 fibril formation support the model of Panel A. Notice that both SegA and SegB A315E fibrils seed LCD monomer (far left and middle left). In contrast, only SegB A315E fibrils seed LCD A315E monomer whereas SegA fibrils barely seed LCD A315E monomer (far right and middle right). Data are shown as mean ± s.d., n=4 independent experiments.

We investigated whether the dagger-and R-shaped folds are relevant to TDP-43 fibrils in human disease, motivated by the irreversibility of fibrils formed by these folds and the likely accessibility to both folds of TDP-CTF. The scarcity of TDP-43 fibrils in autopsied brains of ALS and FTLD patients daunts structure determination of patient-derived TDP-43 fibrils, so we used seeding experiments to examine the possibility that our structures represent folds adopted by TDP-43 in disease. We seeded both SegA and SegB A315E monomer with sarkosyl-insoluble material from brain extracts of two patients with FTLD-TDP, which was previously reported to induce FTLD-TDP pathology in a mouse model^43^. We observe that the sarkosyl-insoluble material was able to seed SegA monomer (Supplementary Figure 1g). This process indicates that TDP-43 in these FTLD-TDP brain extracts contains a well-folded SegA region that can serve as a template for seeded aggregation, thereby providing plausibility for the existence of the dagger-shaped fold in these extracts. In contrast we found that the same brain extracts cannot seed SegB A315E monomer (Supplementary Figure 1g), suggesting that the R-shaped fold may not exist in these extracts. This lack of seeding may mean that the R-shaped fold is specific for patients bearing the TDP-43 A315E mutation. Alternatively, the R-shaped fold may not exist in this particular FTLD patient but may exist in other TDP-43 diseases or disease subtypes such as ALS, especially given that the A315E mutation was discovered in the context of ALS^12^.

We thus propose a one-to-one correspondence between polymorph and disease subtype. More specifically, we speculate that the difference between SegA-sym and SegA-asym are so subtle that they may be associated with the same diseases whereas the differences between the dagger-and R-shaped fold are significant enough to perhaps be associated with different diseases. This hypothesis is supported by : (1) insoluble TDP-43 extracts from different sub-types of FTLD have distinct morphologies and biochemical properties^44^, and (2) fibrils of two peptides whose sequences closely match the dagger-shaped fold and the R-shaped fold (Supplementary Figure 6c) seed cell-expressed TDP-43 into distinct aggregates^41^.

Our finding of polymorphism of TDP-43 fibrils adds TDP-43 to the cohort of other polymorph-forming amyloid proteins^14,28–30^, and raises the question of why polymorphism is common in amyloid fibrils but not in globular proteins. We suggest that the polymorphism observed in irreversible amyloid fibrils arises from a lack of evolutionary pressure to fold into a particular structure that performs an adaptive function. In contrast, each globular protein and functional aggregate has evolved to fold into a single structure with lower free energy than any other structure that its sequence can form. That is, pathogenic amyloid fibrils lack survival advantage and so can adopt multiple conformations that represent different local energy minima in the protein folding landscape. However, a particular polymorph may give rise to a particular disease^45^, perhaps nucleated by a mutation such as TDP-43 A315E or a particular cellular environment as suggested for alpha-synuclein^46^. Polymorph-specific diseases are consistent with results on tau where multiple patients with the same tauopathy all are found to have the same fibril polymorphs and patients with different tauopaties have different fibril polymorphs^29,30,47^.

Our structure of SegB A315E, the first reported cryo-EM fibril structure containing a hereditary mutation, offers a molecular explanation in terms of seeding for cell-to-cell spreading of pathology through the brain^40^. As explained above, our SegB A315E structure indicates that A315E facilitates TDP-43 aggregation through electrostatic attraction with Arg293, and suggests A315T (if the threonine is phosphorylated as speculated) can function in a similar way. As summarized in Figure 4a, wild-type TDP-43 monomer can form either the dagger-shaped or R-shaped fold (or possibly other folds), and both folds can act as seeds to recruit additional monomers into the fibril. In A315E, the inter-layer interaction motif of Arg293-Glu315 provides free positive and negative charges on both ends of any R-shaped fibril that consists of 2 layers or more (Figure 2h), and these free charges can attract additional monomers through long-range electrostatic interactions. Thus the seeding potency of the R-shaped fold with the A315E mutation may exceed that of other folds. Our seeding experiments support this model by showing that SegB A315E fibrils are more effective in seeding TDP-LCD A315E monomer than SegA fibrils (Figure 4b & Supplementary Note 7).

Hereditary mutations have also been proposed to impart pathology by additional mechanisms (Supplementary Note 8), including shifting TDP-43 from a reversible to an irreversible assembly^48,49^. Although we have shown that both dagger-and R-shaped folds form irreversible fibrils, and others have shown that the C-terminal domain can form helical assemblies in liquid droplets^49^, we cannot rule out the possibility that under certain conditions the dagger-and R-shaped folds may also participate in reversible aggregation reported for TDP-43. Energetic analysis suggests this may be possible for the R-shaped fold. The outer two chains in the SegB A315E structure differs noticeably from the inner two chains (Supplementary Note 3), having a lower stabilization energy (0.79 kcal/mol per residue), close to the stabilization energy of the FUS structure (0.66 kcal/mol per residue), which undergoes reversible aggregation, versus the inner chain (0.94 kcal/mol per residue). Conceivably in the native protein, in the absence of the constraint of the Arg293-Glu315 salt bridge and accompanied by certain conformational changes, the R-shaped fold may participate in a reversible fibril (Supplementary Note 9). That is we speculate that the A315E mutation could impose a switch of a reversible to an irreversible fibril.

Our four near-atomic resolution structures of TDP-43 amyloid fibrils establish that: (1) TDP-43 is capable of forming multiple fibrillar structural polymorphs; (2) That two sequence segments of TDP-43 can form distinct stable amyloid cores, one with dagger-shaped folds and the other with R-shaped folds; (3) The R293-E315 salt-bridge in R-shaped fold provides a plausible explanation of enhanced TDP-43 pathology by the ALS related hereditary mutation A315E (and possibly A315Tp); and (4) Energetic analysis highlights the structural features of amyloid fibrils that may lead to both reversible and irreversible aggregation.

## Methods

Methods and materials used in this study are available in supplementary information.

## Acknowledgments

We thank H. Zhou for use of Electron Imaging Center for Nanomachines (EICN) resources. We acknowledge the use of instruments at the EICN supported by UCLA and by instrumentation grants from NIH (1S10RR23057 and 1U24GM116792) and NSF (DBI-1338135 and DMR-1548924). The authors acknowledge NIH AG 054022 and DOE DE-FC02-02ER63421 for support. D.R.B. was supported by the National Science Foundation Graduate Research Fellowship Program.

## Author contributions

Q.C. and D.R.B. designed experiments and performed data analysis. Q.C. expressed and purified constructs, and performed biochemical experiments. Q.C. and D.R.B. prepared cryo-EM samples, and performed cryo-EM data collection and processing. P.G. assisted in cryo-EM data collection and processing. M.R.S. performed solvation energy calculation. All authors analyzed the results and wrote the manuscript. D.S.E. supervised and guided the project.

## Competing interests

D.S.E. is an advisor and equity shareholder in ADRx, Inc.

## Materials & Correspondence

For requests of materials reported in this study, please contact David S. Eisenberg.

## Data and materials availability

All structural data have been deposited into the Worldwide Protein Data Bank (wwPDB) and the Electron Microscopy Data Bank (EMDB) with the following accession codes: SegA-sym (PDB 6N37, EMD-9339), SegA-asym (PDB 6N3B, EMD-9350), SegA-slow (PDB 6N3A, EMID-9349), SegB A315E (PDB 6N3C, EMD-0334). All other data, including the custom software used for solvation energy calculation, are available from the authors upon reasonable request.

## Supplementary Information

### Materials and Methods

#### Construct design

Construct design of TDP-43 segments follows the same strategy reported previously^1^. In addition to TDP-LCD (274-414) reported in our previous study^1^, constructs used in this study include TDP-43 full length (1-414), SegA (311-360), SegB (286-331, wild type or with A315E mutation) and SegAB (286-360). All TDP-43 fragments were conjugated by inserting the cDNA into a pET28a vector that contains an amino terminus conjugation of SUMO protein. The SUMO protein was used to increase solubility and prevent aggregation during expression and purification. The (His)_6_-tag on the amino terminus of the SUMO protein was used for Ni-column purification. The expression sequence of each TDP-43 fragments is as follows:

##### SUMO-TDP-43 (full length, 1-414)

MGSSHHHHHHGSGLVPRGSASMSDSEVNQEAKPEVKPEVKPETHINLKVSDGSSEIFFKIKKTTPLRRLMEAFAKRQGKEMDSLRFLYDGIRIQADQTPEDLDMEDNDIIEAHREQIGGMSEYIRVTEDENDEPIEIPSEDDGTVLLSTVTAQFPGACGLRYRNPVSQCMRGVRLVEGILHAPDAGWGNLVYVVNYPKDNKRKMDETDASSAVKVKRAVQKTSDLIVLGLPWKTTEQDLKEYFSTFGEVLMVQVKKDLKTGHSKGFGFVRFTEYETQVKVMSQRHMIDGRWCDCKLPNSKQSQDEPLRSRKVFVGRCTEDMTEDELREFFSQYGDVMDVFIPKPFRAFAFVTFADDQIAQSLCGEDLIIKGISVHISNAEPKHNSNRQLERSGRFGGNPGGFGNQGGFGNSRGGGAGLGNNQGSNMGGGMNFGAFSINPAMMAAAQAALQSSWGMMGMLASQQNQSGPSGNNQNQGNMQREPNQAFGSGNNSYSGSNSGAAIGWGSASNAGSGSGFNGGFGSSMDSKSSGWGM

##### SUMO-SegA (311-360)

MGSSHHHHHHGSGLVPRGSASMSDSEVNQEAKPEVKPEVKPETHINLKVSDGSSEIFFKIKKTTPLRRLMEAFAKRQGKEMDSLRFLYDGIRIQADQTPEDLDMEDNDIIEAHREQIGGMNFGAFSINPAMMAAAQAALQSSWGMMGMLASQQNQSGPSGNNQNQGNMQ

##### SUMO-SegB (286-331)

MGSSHHHHHHGSGLVPRGSASMSDSEVNQEAKPEVKPEVKPETHINLKVSDGSSEIFFKIKKTTPLRRLMEAFAKRQGKEMDSLRFLYDGIRIQADQTPEDLDMEDNDIIEAHREQIGGQGGFGNSRGGGAGLGNNQGSNMGGGMNFGAFSINPAMMAAAQAALQ

##### SUMO-SegB A315E (286-331)

MGSSHHHHHHGSGLVPRGSASMSDSEVNQEAKPEVKPEVKPETHINLKVSDGSSEIFFKIKKTTPLRRLMEAFAKRQGKEMDSLRFLYDGIRIQADQTPEDLDMEDNDIIEAHREQIGGQGGFGNSRGGGAGLGNNQGSNMGGGMNFGEFSINPAMMAAAQAALQ

##### SUMO-SegAB (286-360)

MGSSHHHHHHGSGLVPRGSASMSDSEVNQEAKPEVKPEVKPETHINLKVSDGSSEIFFKIKKTTPLRRLMEAFAKRQGKEMDSLRFLYDGIRIQADQTPEDLDMEDNDIIEAHREQIGGQGGFGNSRGGGAGLGNNQGSNMGGGMNFGAFSINPAMMAAAQAALQSSWGMMGMLASQQNQSGPSGNNQNQGNMQ

#### Protein purification and SUMO-tag cleavage

Protein purification follows the same protocol reported previously^1^. Briefly, all SUMO-tagged proteins were expressed in *Escherichia coli* BL21 (DE3) strain. LB media with 50 µg/ml kanamycin was used for cell culture. After bacterial cells were cultured at 37 °C to an OD600 of 0.6-0.8, 1 mM isopropyl β-D-1-thiogalactopyranoside (IPTG) was added for protein expression and the cells were further cultured at 25 °C for 3 h. Cells were harvested and resuspended in 20 mM Tris-HCl, pH 8.0, 500 mM NaCl, 20 mM imidazole, 10% (v/v) glycerol, supplemented with 1% (v/v) Halt Protease Inhibitor single-use cocktail (Thermo Scientific), and were sonicated (3s on/3s of cycle, 10 min) and centrifuged (24,000 g for 20 min) to get cell lysate. The cell lysate was mixed with homemade NucA nuclease (5000 U per liter of cell culture) and filtered before loading onto a HisTrap HP column (GE healthcare) for purification. The HisTrap column was pre-equilibrated with 20 mM Tris-HCl, pH 8.0, 500 mM NaCl and 20 mM imidazole, and sample was washed with 20 mM Tris-HCl, pH 8.0, 500 mM NaCl and 200 mM imidazole and eluted with 20 mM Tris-HCl, pH 8.0, 500 mM NaCl and 500 mM imidazole. Eluted protein was concentrated using Amicon Ultra-15 centrifugal filters (Millipore) and stored at −80 °C for future use.

In order to form fibrils of the TDP-43 fragments, the SUMO-tag of each fragment was removed before aggregation. SUMO-tagged protein was mixed with 100:1 (weight basis) homemade ULP1 protease, and incubated for 1 hour. When analyzed by SDS-PAGE, samples mixed with ULP1 at 0 hour show bands of intact SUMO-tagged protein; after 1 hour of cleavage, bands of free SUMO and TDP-43 fragments were shown on the gel, as exampled by SegA and SegB A315E cleavage shown in Supplementary Figure 1a.

#### Fibril preparation and optimization

SUMO-tag removed proteins were used for fibril preparation. For all TDP-43 fragments, the starting fibril preparation conditions are the same. Protein was diluted to 50 µM concentration with 20 mM Tris-HCl, pH 8.0, 150 mM NaCl, 10 µM DTT and filtered using 0.1 µm Ultrafree-MC-VV centrifugal filters (Millipore), and incubated at 37 °C with shaking for 3 days. The morphology of each sample was checked by negative stain TEM.

In order to achieve the necessary fibril concentration and distribution for cryo-EM structure determination, the fibril growth condition was optimized for each sample. The optimization includes determining the appropriate concentration of proteins (final values ranging from 3 µM to 100 µM monomer concentration), incubating the solution without shaking, incubation temperature (4 °C or 25 °C or 37 °C), incubation time (overnight to more than 2 weeks), buffer (Tris or phosphate), salt concentration (50 mM to 500 mM NaCl) and seeding with pre-formed fibrils. After optimization of all TDP-43 fragments, SegA and SegB A315E were found to have the best fibril morphology, distribution, and concentration and were therefore selected for cryo-EM structure determination. The optimized fibril morphology of each fragment is shown in Supplementary Figure 1b.

The optimized fibril growth condition for SegA was 50 µM protein concentration, incubation in 20 mM Tris-HCl, pH 7.4, 500 mM NaCl, 10 µM DTT at 37 °C with 3 days of shaking. Two percent (molar ratio, monomer equivalent) pre-formed SegA fibrils were added as seeds. After shaking, SegA fibrils were collected by centrifugation at 9,000 g for 5 min and washed twice and resuspended to the starting volume with the identical buffer as above, except using 150 mM NaCl instead of 500 mM. Fibrils in resuspension buffer were further incubated without shaking for 3 days at room temperature. Before cryo-EM sample preparation, SegA fibrils were again collected by centrifugation, and re-suspended with 150 mM NaCl buffer at 5% volume of previous incubation (20 × concentrating), and the concentrated solution was heated to 75 °C for 30 min with a PCR machine. The final heating step was necessary to de-clump SegA fibrils and optimize fibril distribution.

The optimized fibril growth condition for SegB A315E was 50 µM initial protein concentration, and incubation in 20 mM Tris-HCl, pH 8.0, 150 mM NaCl, 10 µM DTT, at 37 °C without shaking. Two percent (molar ratio, monomer equivalent) pre-formed SegB A315E fibrils were added as seeds. After incubation, SegB A315E fibrils were collected by centrifugation at 9,000 g for 5 min and washed twice by the same incubation buffer at 5% volume of previous incubation (20 × concentrating), and the concentrated solution was used for cryo-EM sample preparation.

#### Negative stain transmission electron microscopy (TEM)

Negative stain TEM samples were prepared by applying 5 µl of fibril solution to glow-discharged 400 mesh carbon-coated formvar support films mounted on copper grids (Ted Pella, Inc.) and incubating on the grid for 2 min. The samples were then blotted off and the grids were stained with 3 µl of 2% uranyl acetate for 1 min. The grids were washed with an additional 3 µl of 2% uranyl acetate and allowed to dry for 1 min. Each grid was imaged using a T12 (FEI) electron microscope.

#### Cryo-EM data collection and processing

2 µl of fibril solution was applied to a glow-discharged Quantifoil 1.2/1.3 electron microscope grid and plunge-frozen into liquid ethane using a Vitrobot Mark IV (FEI). Data were collected on a Titan Krios (FEI) microscope equipped with a Gatan Quantum LS/K2 Summit direct electron detection camera (operated with 300 kV acceleration voltage and slit width of 20 eV). Super-resolution movies were collected on a Gatan K2 Summit direct electron detector with a nominal physical pixel size of 1.07 Å /pixel (0.535 Å /pixel in super-resolution movie frames) with a dose per frame of ∼1.1 e^-^/Å ^2^. A total of 40 frames with a frame rate of 5 Hz were taken for each movie resulting in a final dose of ∼44 e^-^/Å ^2^ per image. Automated data collection was driven by the Leginon automation software package^2^.

Micrographs containing crystalline ice were used to estimate the anisotropic magnification distortion using mag_distortion_estimate^3^. CTF estimation was performed using CTFFIND 4.1.8 on movie stacks with a grouping of 3 frames and correction for anisotropic magnification distortion^4^. Unblur^5^ was used to correct beam-induced motion with dose weighting and anisotropic magnification correction, resulting in a physical pixel size of 1.06 Å /pixel and 1.064 Å /pixel for the SegB and SegA data sets, respectively.

All particle picking was performed manually with SegB A315E data set fibrils picked as a group and SegA data set fibrils were picked separately in groups of slow twister and fast twisters using EMAN2 *e2helixboxer.py*^6^ (we could not discriminate between SegA-sym and SegA-asym by eye due to their similar crossover distances and morphology; therefore, SegA-sym and SegA-asym particles were later separated by 2D classification and 3D classification). Particles were extracted in RELION using the 90% overlap scheme into 288 pixel boxes for SegA and 320 pixel boxes for SegB A315E. Classification, helical reconstruction, and 3D refinement were used in RELION as described^7–9^. In general, we performed 2D Classifications with higher tau_fudge (ranging from values of 4 to 12) to identify, and select for future 3D classification, particles contributing to 2D classes showing β-strand separation along the helical axis. We performed 3D classification with the estimated helical parameters from each structure and an elongated Gaussian blob as an initial model to generate starting reconstructions. We ran additional 3D classifications using the preliminary reconstructions from the previous step to select for particles contributing to homogenous classes (stable helicity and separation of β-strands in the X-Y plane). Typically, we performed Class3D jobs with K=3 and manual control of the tau_fudge factor and healpix to reach a resolution of ∼5-6 Å to select for particles that contributed to the highest resolution class for each structure. For the SegB and SegA-slow reconstructions, we performed high-resolution gold-standard refinement as described^9^. For SegA-sym and SegA-asym, perhaps due to low particle numbers, Refine3D seemed unable to adequately regulate working resolution to refine with high enough resolution factors to allow SegA-sym and SegA-asym reconstructions to refine past 4.8 Å, and gold-standard refinement did not result in near-atomic resolution maps. Therefore, we employed a “sub-optimal” manual refinement protocol using all particles against a single reference map as described^7,10^. Final overall resolution estimates were calculated from the corrected Fourier shell correlations of two half maps, using high-resolution phase randomization as implemented in RELION PostProcess to account for effects of a soft-edged solvent mask and to monitor possible overfitting for the SegA-sym and SegA-asym manual refinements^11^. For the SegB and SegA-slow gold-standard refinements, we report the 0.143 FSC resolution cutoff; however, for the SegA-sym and SegA-asym “sub-optimal” refinements we report the 0.5 FSC resolution cutoff (the particles contributing to the final map were split *a posteriori* into even and odd micrographs to prevent particles from the same fibril contributing to different reconstructions and two half-maps were reconstructed using the fully refined center and orientation parameters)^11^. In all cases except the SegA-slow, the default value of 30% was used for the *helical_z_percentage* during 3D classification and refinement. Since the crossover distance of the SegA-slow varied from 1200-2000 Å, we used 10% for the *helical_z_percentage* to minimize the blurring due to the variable twist as described^12^. Reconstructions were compared against the reference-free 2D class averages to confirm their validity. In addition, *de novo* model-building also confirms the validity of the 3D reconstructions.

Specific determination of helical symmetry for all structures was performed as follows. For SegA-sym, SegA-asym, and SegB A315E, we estimated the crossover distances using a combination of manual inspection of the micrographs and inspection of 2D class averages using larger particle box sizes to capture the entire crossover distance (box sizes from 432 to 1024 pixels were tested). In the case of the SegA-sym and the SegB A315E, the mirror symmetry axis parallel to the helical axis in the 2D class averages indicated either a two-start helix symmetry with a rise of 2.4 Å or a C_n_ rotational symmetry. Computed diffraction patterns from the 2D class averages indicated there was no meridional reflection at 4.8 Å, indicating the lack of C_n_ rotational symmetry. Therefore, for both of these species, a helical rise of 2.4 Å was assumed and the helical twist was calculated from the previously estimated crossover distance. Convergence to a stable helicity and high-resolution refinement to near-atomic resolution allowing *de novo* model building confirmed the helical symmetry. For SegA-slow, the entire crossover distance could not be captured by the 1024 box size, so the crossover distance was estimated from a combination of manual inspection of micrographs and a “stitched” together fibril using the 2D classes from the 288 box size (Supplementary Figure 3f). Class averages, as well as the computed diffraction pattern from the “stitched” fibril, confirm the presence of a meridional reflection at 4.8 Å, indicating C_n_ rotational symmetry. Therefore, we assumed a helical rise of 4.8 Å and we calculated the helical twist from the previously estimated crossover distance. High-resolution refinement to near-atomic resolution allowed *de novo* model building, confirming the helical symmetry. In addition, reconstructions with a helical rise of 2.4 Å did not show clear side chain density, and projections of the reconstruction did not match the 2D class averages (Supplementary Figure 3e). For SegA-asym, the crossover distance was estimated from a combination of inspection of the micrographs, and 2D class averages with larger box sizes. Due to the lack of apparent two-fold symmetry in the 2D class averages, we assumed a helical rise of 4.8 Å and calculated the helical twist from the measured cross-over distances. Refinement to near-atomic resolution demonstrate that the two SegA-asym monomers within each asymmetric unit are not identical, confirming the lack of two-fold symmetry.

#### Atomic model building

We sharpened the refined maps using phenix.auto_sharpen^13^ at the resolution cutoff indicated by half-map FSC, and subsequently built atomic models into the refined maps with COOT^14^. For the three SegA polymorphs, we built the model of the slow twister first owing to its high resolution (3.3 Å). The density map of the slow twister clearly shows a cyclic mainchain with the amino and carboxy termini at the center of the fibril. The initial model was built with both N-C orientations, and only the orientation shown in the current model fit all the side chain densities. The conservation of the dagger-fold in the maps of the fast symmetric and fast asymmetric twister, especially the signature ^334^WGMMGML^340^ motif, allowed us to rigid-body fit the model of the slow twister into the fast symmetric and fast asymmetric maps. Following rigid-body fitting, we manually inspected and adjusted each residue.

After building atomic models for the main chain of SegA polymorphs, we observed extra density peripheral to the dagger-shaped fold around Gln331 of SegA-slow and Met323 of both SegA-asym and SegA-slow (Figure 1c-d). We believe this density arises from unstable interactions (either due to weak binding or binding of a mixture of different residues) of other protomers with the core protofilaments. We could not build a definitive model due to ambiguity of the side-chain densities; however, we built a hypothetical model into the density near Gln331 of SegA-slow based on the speculation that this density represents part of the continuous dimer interface found in SegA-sym and SegA-asym.

During model building of SegA-slow, we found that the model that fully recapitulates the registration of the fast symmetric twister was not favorable due to the steric clash at L340 and unexplained side chain density at G338 (Supplementary Figure 5d). Therefore, we built the model with a registration that has a 2 residue offset compared to the fast symmetric twister. We believe that this registration shift is allowed by the similarity of nearby residues (Gln331 from dagger-fold flanked by Ser342 and Gln344 from the short chain, compared to Gln331 flanked by Gln344 and Gln346), and is caused by the lack of the hydrophobic anchor formed by Met336 and Leu340 (Supplementary Figure 5e). A similar registration shift was observed in previous reported crystal structures of steric-zippers and was termed registration polymorphism^15^. The extra densities near Met323 in SegA-asym and SegA-slow could represent an additional dimer interface that protects Ala321, Met323 and Ala325 from the solvent, but without unambiguous side chain density, we could not build in any models for these densities.

The model building of SegB A315E was initiated by the inspection of the density map near the homodimer interface on the neck of the inner-chain R-shaped fold. The side chain density and nearby environment indicates the atomic model of this region should feature 4-5 small side chain residues flanked by a big side chain residue at each side, and the only motif in the SegB A315E sequence that matches this criterion is ^293^RGGGAGL^299^, with only one orientation feasible considering the length of density at each side of this motif. With the model of ^293^RGGGAGL^299^ built, the rest of model was built by adding the residues one-by-one. All sidechains fit the density unambiguously. The observation that Glu315 forms a salt-bridge with Arg293 further supports this model. After we built the inner R-shape fold, we speculated that the outer R-shape fold was not a continuation of the same chain as the inner R-shape fold because there were not enough residues in the sequence to accommodate the outer R-shape fold. Therefore, the model of the outer R-shape fold was built *de novo* similarly to the inner-R-shape fold.

For all structures, after we built the initial model, we used in-house scripts to generate a 5-layer model using the helical symmetry of the refined map, and used phenix.real_space_refine for model refinement^16^. After several rounds of refinement, we adjusted the orientation of the main chain oxygen and nitrogen to facilitate main chain hydrogen bond within the β-sheet, and we applied these mainchain hydrogen bond restraint for further real-space refinement. As the last step, the rotamer of each serine, glutamine and asparagine residue was manually inspected to ensure the energy favorable hydrogen binding. The final model was validated by MolProbity^17^.

The G304W model of SegB A315E shown in Supplementary Figure 6b was generated by manually changing Gly304 residues of 1-layer SegB A315E model (four glycine residues in four chains) to tryptophan in COOT, and we generated a 5-layer mutated model using the same script as for original SegB A315E model. We refined this 5-layer mutated model by phenix.real_space_refine using the same density map we used to refine the original model. The refined model shows no Ramachandran angles and rotamer outliers, and no significant clashes. The statistics of SegA A315E G304W model are shown in Supplementary Table 3.

#### Energetic calculation

The stabilization energy is an adaptation of the solvation free energy described previously^18^, in which the energy is calculated as the sum of products of the area buried of each atom and its corresponding atomic solvation parameter (ASP). ASPs were taken from our previous work^18^. Area buried is calculated as the difference in solvent accessible surface area (SASA) of the reference state (i.e. unfolded state) and the SASA of the folded state. The reference state was measured absent all other atoms in the structure but the residue “i” and main chain atoms of residue i-1 and i+1. The SASA of the folded state was measured for each atom in the context of all amyloid fibril atoms. Fibril coordinates were extended by symmetry by three to five chains on either side of the reported molecule, to ensure the energetic calculations were representative of the majority of molecules in a fibril, rather than a fibril end. To account for energetic stabilization of main chain hydrogen bonds, the ASP for backbone N/O elements was reassigned from −9 to 0 if they participated in a hydrogen bond. Similarly, if an asparagine or glutamine side chain participated in a polar ladder (two hydrogen bonds per amide), and was shielded from solvent (SASA_folded_ < 5 Å^2^), the ASPs of the side chain N and O elements were reassigned from −9 to 0. Lastly, the ASP of ionizable atoms (e.g. Asp, Glu, Lys, His, Arg, N-terminal amine, or C-terminal carboxylate) were assigned the charged value (−37/−38) unless the atoms participated in a buried ion pair, defined as a pair of complementary ionizable atoms within 4.2 Å distance of each other, each with SASA_folded_ < 40 Å^2^). In that case, the ASP of the ion pair was reassigned to −9. In the energy diagrams, a single color is assigned to each residue, rather than each atom. The color corresponds to the sum of solvation free energy values of each of the atoms in the residue. The energy reported for FUS in Figure 3 is the average over 20 NMR models. The standard deviation is 1.8 kcal/mol.

#### Thermo-stability assays of TDP-43 and FUS fibrils

To validate the energetic calculations and test the stability of TDP-43 segment fibrils experimentally, pre-formed TDP-43 segment fibrils were heated at 75 °C for 30 min in a PCR machine, and the existence of the fibrils were checked by negative stain EM. All fibrils were heated in 20 mM Tris pH 7.4, 150 mM NaCl, 10 µM DTT at 50 µM monomer-equivalent concentration.

FUS protein was purified and FUS fibrils was formed the same way as previously reported [ref]. FUS protein was expressed using pHis-parallel-mCherry-1FUS214 plasmid that contain the low complexity domain of FUS (1-214) conjugated to mCherry. Purification was done by Ni column and size exclusion column. The hydrogel of FUS was formed by incubating purified mCherry-FUS at 4 °C for 2 weeks at a concentration of 1 M. After hydrogel formation, a scrape of hydrogel was mixed with 20 mM Tris pH 7.4, 150 mM NaCl, 10 µM DTT to approximately 20x of its original volume (∼50 µM monomer concentration). The mixture was allocated to three PCR tubes, one without heat and two heated at 60 °C and 75 °C for 30 min, respectively. All three samples were checked by negative stain EM for fibrils. The sample without heat contains abundant fibrils on the EM grid and the sample with heat contains no visible fibrils anywhere on the EM grid.

#### ThT assays

Protein was diluted into 12 µM in 20 mM Tris-HCl, pH 7.4, 500 mM NaCl, 10 µM DTT, 30 µM ThT, and filtered using 0.1 µm Ultrafree-MC-VV centrifugal filters (Millipore). Filtered solution was mixed with or without 100:1 (weight basis) ULP1, with or without 5% (molar ratio, monomer equivalent) pre-formed fibril seeds (or 1% v/v for brain extract seeds), and was pipetted into a polybase black 384-well plate with optical bottom (Thermo Scientific) and incubated at 37 °C without shaking. ThT fluorescence was measured with excitation and emission wavelength of 440 and 480 nm, respectively, using FLUOstar Omega plate reader (BMG LABTECH). The aggregation curves were averaged from four independent measured replicates and error bars show s.d. of replicate measurements. Fibril seeds were sonicated (1s on/1s off cycle, 3 min) before use to increase the seeding ability.

The brain extracted materials were a generous gift from Prof. Virginia Lee’s lab. Brain extract #1 was from a patient diagnosed as FTLD-TDP, whereas brain extract #2 was from a patient diagnosed as FTLD-TDP-GRN. Both brain extracts were extracted from the frontal cortex, and estimated TDP-43 concentration were ∼500 ng/ml (measured by ELISA assays) and estimated total protein concentration were 2.5 mg/ml for brain extract #1 and 4 mg/ml for brain extract #2 (measured by BCA assays).

The samples mixed with pre-formed fibril seeds had an additional ThT background reading because of the seeds. To better compare the curves of samples with and without seeding and estimate the ThT signal change that represented the formation of seeded fibrils, we removed the background by subtracting the value of the first reading from each curve. To normalize the different ranges of fluorescence readings observed from each experiments (probably due to the different fluorescence gain settings of the plate reader), we normalized the readings to make the minimum mean value in each panel 0%, and the maximum mean value in each panel 100%.

#### TDP-CTF aggregation assays

The aggregation assays of pathological fragments of TDP-43 (TDP-CTF) were performed largely the same as previous reported^1^. The SUMO-TDP-CTF wild type and mutants were diluted into 0.2 mg/ml with 20 mM Tris-HCl, pH 8.0, 150 mM NaCl, 10 µM DTT and filtered by 0.1 µm Ultrafree-MC-VV centrifugal filters (Millipore). To simplify the experiment, we did not remove SUMO-tag by ULP1 protease because we found that wild type TDP-CTF aggregates after overnight incubation even with the protection of SUMO. Filtered samples were incubated at 4 °C overnight, and separated into supernatant and pellet fractions by centrifugation at 18,000 g for 3min at 4 °C. The pellet was resuspended with the same buffer and volume, and both supernatant and pellet were mixed with 3:1 (v/v) of NuPAGE LDS sample buffer (Invitrogen) and heated to 100 °C for 10 min. The samples before incubation were also prepared in the same way. All samples were separated by NuPAGE 4-12% Bis-Tris gels (Invitrogen) and stained by Coomassie blue. TDP-CTF A324W that had been shown to reduce TDP-CTF aggregation previously^1^ was also tested in this study as a control to make sure that we can get the similar results with or without ULP1 cleavage.

### Supplementary Text

#### Note 1 (Detailed comparison of the dagger-shaped fold in three SegA polymorphs)

All three polymorphs feature a largely conserved dagger-shaped fold (Figure 2a&b), with the hydrophobic residues from Phe313 to Ala341 forming tight hydrophobic interactions (Figure 2c). Detailed single chain superposition shows that SegA-sym and the long chain of SegA-asym have nearly identical dagger-shaped folds, with the “stem” of the long chain SegA-asym (Ala341 to Ser347) bent approximately 40° (Figure 2c). This bending of the long chain SegA-asym stem forces a 40° bend in the “blade” of the SegA-asym short chain relative to the reference SegA-sym fold, corresponding with the shift in registration of the stem hydrophobic interdigitation (Supplementary Figure 5a, left). The blade region of the SegA-asym short chain (Pro320 to Met337) is still conserved (Supplementary Figure 5a, middle). In addition, the dagger-shaped fold in SegA-slow is also largely conserved; however, Ala341 to Ser347 of the SegA-slow stem are bent approximately 50° compared to SegA-sym (Supplementary Figure 5a, right), causing a significant conformational change at Met336. Further, the interaction of the termini in SegA-slow causes a conformational change at Phe313, Phe316 and Met339, and M311 is now visualized and participating in the hydrophobic core (Supplementary Figure 5a, right). We note that the dagger tip (residue 320-334) is almost identical in all four dagger-shaped fold of three SegA polymorphs despite differences in local environment. This high degree of structural conservation suggests this core region of dagger-shaped fold is thermodynamically stable, kinetically accessible and is likely to be present in other polymorphs (perhaps including disease relevant TDP-43 aggregates).

#### Note 2 (Detailed comparison of dimer interfaces in three SegA polymorphs)

The protofilament interface in SegA-sym and SegA-asym are similar (Supplementary Figure 5b&c) with slight differences in the region that have 40° bend (Supplementary Figure 5c, right). We name this the continuous interface since it is formed by two continuous chains, compared to the broken interface in SegA-slow formed by the interaction of the termini (Supplementary Figure 5a). The continuous interface has a hydrophobic core formed by Met336 and Leu340 from both monomers in the middle and a hydrogen bond network towards the edges (Figure 2d & Supplementary 5b). The broken interface is formed largely by the tight packing of Met359 and the hydrogen bonding network of the peripheral asparagine residues (Supplementary Figure 5b). We observed only part of the continuous interface in SegA-slow, probably due to the conformational change of Met336 (Supplementary Figure 5d&e, see model building in Methods for details).

#### Note 3 (information provided by SegA-slow)

The close contact between the N-and C-terminal in the long chain of SegA-slow suggests that SegA-slow is not compatible with any TDP-43 segments that are longer than SegA, so that this structure is probably a truncation artifact. However, we still believe this structure provides valuable information that helps us understand the aggregation of TDP-43. First, the SegA-slow structure contains the dagger-shaped fold, demonstrating its conservation in the three different polymorphs we observe for SegA (Supplementary Figure 5a). We note that the incompatibility of SegA-slow to full-length TDP-43 is coming from the tail region (residues 347-360) that is extra to the dagger-shaped fold (Supplementary Figure 8a, upper panel). Thus, taken alone, the dagger-shaped fold in SegA-slow (residues 312-346) is still compatible with full-length TDP-43, as in SegA-sym and SegA-asym. Second, the tail region hairpin structure (residues 342-360) in SegA-slow is also compatible with full-length TDP-43 when separated from the dagger-shaped fold, with free N-and C-terminals (Supplementary Figure 8a bottom panel). This hairpin structure is formed by tightly packed hydrophilic zippers stabilized with extensive hydrogen bonds (Supplementary Figure 8b). Based on the analysis above, we speculate that this hairpin structure may also be involved in TDP-43 aggregation as another segmental polymorph. Further study is needed to test our hypothesis as well as identify whether this hairpin contribute to reversible or irreversible aggregation of TDP-43. Third, our SegA-slow structure may provide, to our knowledge, the first atomic evidence of secondary seeding of amyloid fibrils. Secondary seeding is considered as seeding off the side of pre-formed fibrils, in contrast to primary seeding where molecules add to the top or bottom of pre-formed fibrils. In this case, the extra density near Q331 and M323 of the long chain of SegA-slow indicates that the secondary seeding can occur on these regions. Because the long chain of SegA-slow contain all 50 residues in SegA, the extra density must come from other SegA molecules, therefore supporting a secondary seeding mechanism. In comparison, similar extra density has been observed in other amyloid fibril structures, such as PHF tau^12^; but in these structures the ordered fibril core does not contain all residues of sample molecule, so that we cannot be sure that these densities are coming from a different chain as examples of secondary seeding, or from flexible residues from the same molecule outside the fibril core.

#### Note 4 (Structural analysis of SegB A315E)

The overall R-shaped folds of the inner and outer chain of the SegB A315E fibril are largely similar with some salient differences. The head region of the R-fold inner chain is formed by the hydrophobic interaction of Leu299, Met307, Met311 and Phe313, as well as by Asn302 and Ser305 hydrogen bonding; whereas in the outer chain, Leu299 is flipped outside the head to mediate the heterodimer interaction between the inner and outer chain (Figure 2f). Consequently, the head region of the outer chain is formed by the hydrophobic interaction of Met307, Met311 and Phe313, together with hydrogen bonds formed by Asn301, Asn302 and Ser305 (Figure 2f & Supplementary Figure 5f). The neck region of both chains is formed by hydrophobic interaction between Ala297 and Phe313, the Arg293-Glu315 salt-bridge. The 70° bend of the Phe316-Asn319 leg of the outer chain relative to the inner chain is probably caused by the heterodimer interface formed by the legs of the inner and outer chain that restricts the conformation of inner chain.

#### Note 5 (Structure analysis of SegB A315E dimer interfaces)

The homodimeric interfaces (between two pseudo-2_1_ axis related inner chains) are formed by small side chain residues (Gly298 and Gly300) in the middle (reminiscent of the homodimer interface in the fibril structure of PHF of tau from Alzheimer’s disease^12^) and hydrogen bonds among Gln303, Gly295, Ala297 and Gln301 (Figure 2g, left and Supplementary Figure 5g). The heterodimeric interfaces (between inner chain and outer chain) are formed by hydrophobic interaction of Ile318, Phe316 with Phe289, the insertion of Leu299 into the pocket formed by the main chain of Asn306-Gly309, hydrophobic interaction of Phe289 (from another inner chain) and the main chain Asn303-Gly304, and hydrogen bonds among Asn312, Gly290, Ser292 and Met311 (Figure 2g, right and Supplementary Figure 5g).

#### Note 6 (Structure analysis of mutagenesis studies)

In SegA-sym, three of these mutation sites (Ala324, Leu330 and Met337) point toward the center of the dagger-shaped fold and the other one (Gln331) points toward the “continuous dimer interface” (Figure 2b & Supplementary Figure 6a), and all three are buried in a tightly packed interface that cannot tolerate a tryptophan replacement. In contrast, Ala326 points away from the dagger-shaped fold and is solvent exposed, so that a tryptophan mutation is allowed.

In SegB A315E, Gln303 of the inner chain is buried within the inter-chain interface whereas Gly304 of both inner and outer chains are located in a less tightly packed environment, allowing the tryptophan mutation at position 304 (Figure 3a & Supplementary Figure 6b). Meanwhile, R293E/A315E double mutation is expected to disrupt the R-shaped fold by creating electrostatic hindrance, whereas R293E/A315R should restore the salt-bridge of Arg293 and Glu315 and facilitate the formation of the R-shaped fold.

#### Note 7 (Analysis of ThT seeding assays)

The observation that both SegA and SegB 315E fibrils can seed LCD monomer (Figure 4b, far-left and middle-left panel), indicate that the LCD can adopt both the dagger-and R-shaped fold when seeded and also suggest that roughly equal seeding ability of both folds. Meanwhile, we found SegA fibrils barely seed LCD A315E monomer (Figure 4b, middle-right panel) whereas SegB A315E fibrils can seed LCD A315E monomer (Figure 4b, far-right panel). We believe these results are unlikely to come from the incompatibility of the dagger-shaped fold with A315E mutation because the Ala315 in the dagger-shaped fold is facing towards the solvent so that A315E mutation should be allowed. Instead, we believe the results shown in Figure 4b middle-right and far-right panels indicate that the seeding ability of the R-shaped fold is much higher than the dagger-shaped fold when seeding A315E monomer. In this case, the observation that SegA fibrils barely seed LCD A315E monomer can be explained by the enhanced primary nucleation and self-seeding of LCD A315E monomer via the R293-E315 salt-bridge, so that the self-seeded aggregation overwhelms the seeding of SegA fibrils. Taken together, we think that the seeding experiments shown in Figure 4b support our hypothesis that the salt-bridge introduced by A315E mutation accelerates the aggregation of the R-shaped fibrils.

#### Note 8 (Possible mechanisms of other hereditary mutations)

The structures of the dagger-and R-shaped folds may help explain the role of other hereditary mutations. For instance, if the R-shaped fold does indeed participate in reversible aggregation, mutations that disrupt it could promote a transition to a more irreversible amyloid core, such as the dagger-shaped fold, as previously hypothesized^19^ (Supplementary Figure 6d). On the other hand, we observe that many hereditary mutations would have a neutral, or even deleterious, effect on the dagger-and R-shaped folds. This is consistent with the idea that there are other observed mechanisms for mutation-enhanced pathology, such as toxic gain-of-function RNA splicing activity^20^. In addition, these mutations may influence the formation of other amyloid cores not discovered here.

#### Note 9 (Possible conformational change of the R-shaped fold in reversible aggregates of TDP-43)

We speculate that large structural and energetic differences between the outer and inner chains is perhaps due to the flexibility of the R-shaped fold owing to its high glycine content and relatively small hydrophobic core. In humans lacking the A315E mutation, the absence of the Arg293-Glu315 salt bridge would eliminate the electrostatic attraction that helps to fix the R-shape, perhaps allowing the molecule to partially unfold to a more open U-shape. This conformational change may further compromise the stability of the R-shaped fold by promoting the disassembly of the four-protofibril bundle given the observation that the homodimer interface of the R-shaped fold is relatively weak (Figure 2g) and formation of the heterodimer interface requires the closed conformation of the inner R-shaped folds. Therefore, the transition from reversible to irreversible assemblies may occur through a U-R switch where a more open and less bundled U-shape fold converts to a more compact and more bundled R-shape. The U-R switch is possible with both wild-type and A315E, although perhaps the wild-type sequence would have a higher transition state energy owing to the loss of the Arg293-Glu315 interaction. We note that our energy analysis relies on the structure of one reversible fibril and that we have no experimental evidence regarding the role of the R-shape in reversible TDP-43 aggregation; therefore, further research identifying structures of reversible fibrils and interrogating the role of the R-shape in TDP-43 reversible aggregation is needed.

## Supplemental Figures

**Supplementary Figure 1a-d.**
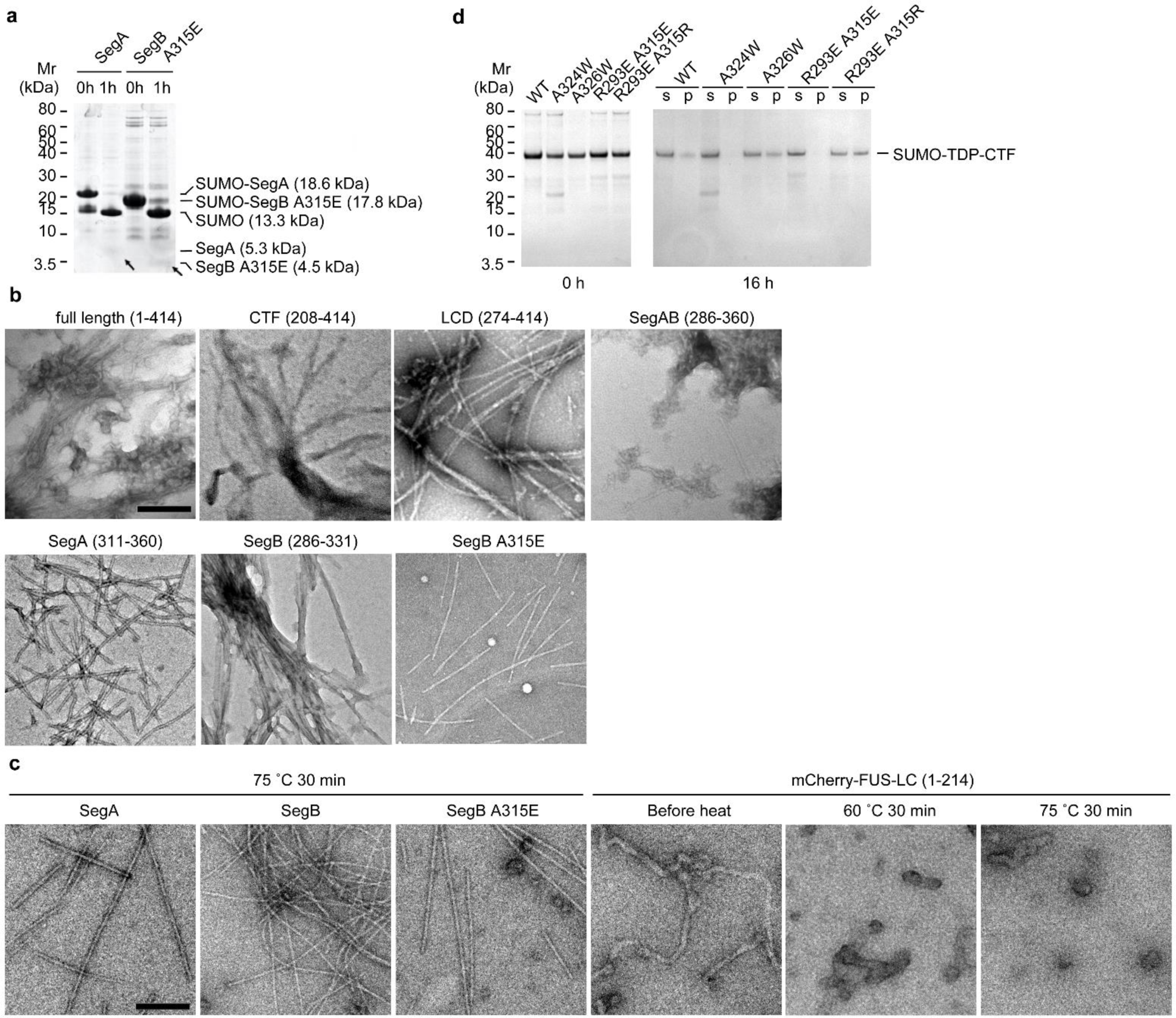
Biochemical characterization of TDP-43 segments. **a**, SDS-PAGE shows the cleavage of SegA and SegB A315E. SUMO-tagged segments are purified with Ni-column, and mixed with protease UPL1 to remove SUMO tag (0h). After 1 hour of cleavage (1h), bands corresponding to SUMO tag alone and TDP-43 segments (indicated by arrows) are shown on the gel, indicating successful cleavage. **b & c**, Negative stain electron microscope (EM) images of fibrils formed by TDP-43 segments (b) without heating or (c) heated before EM sample preparation. Fibrils of mCherry-FUS-LCD with or without heating were also observed with EM as a control. (Scale bar = 200 nm) Notice that the amount of fibrils in each image does not necessary corresponding to the total amount of fibrils in each sample, since the distribution of fibrils on EM grids was not even, especially for clumped fibrils. **d**, Aggregation assays of the pathological fragment TDP-CTF (208-414). Wild type (WT) and mutant TDP-CTFs conjugated with SUMO tags were incubated at 4 °C overnight. Samples were separated into supernatant (s) and pellet (p) by centrifugation and analyzed by SDS-PAGE. Notice that TDP-CTF A326W and R293E-A315R behave similarly as TDP-CTF WT, whereas the other two mutants show reduced aggregation.

**Supplementary Figure 1e-g.**
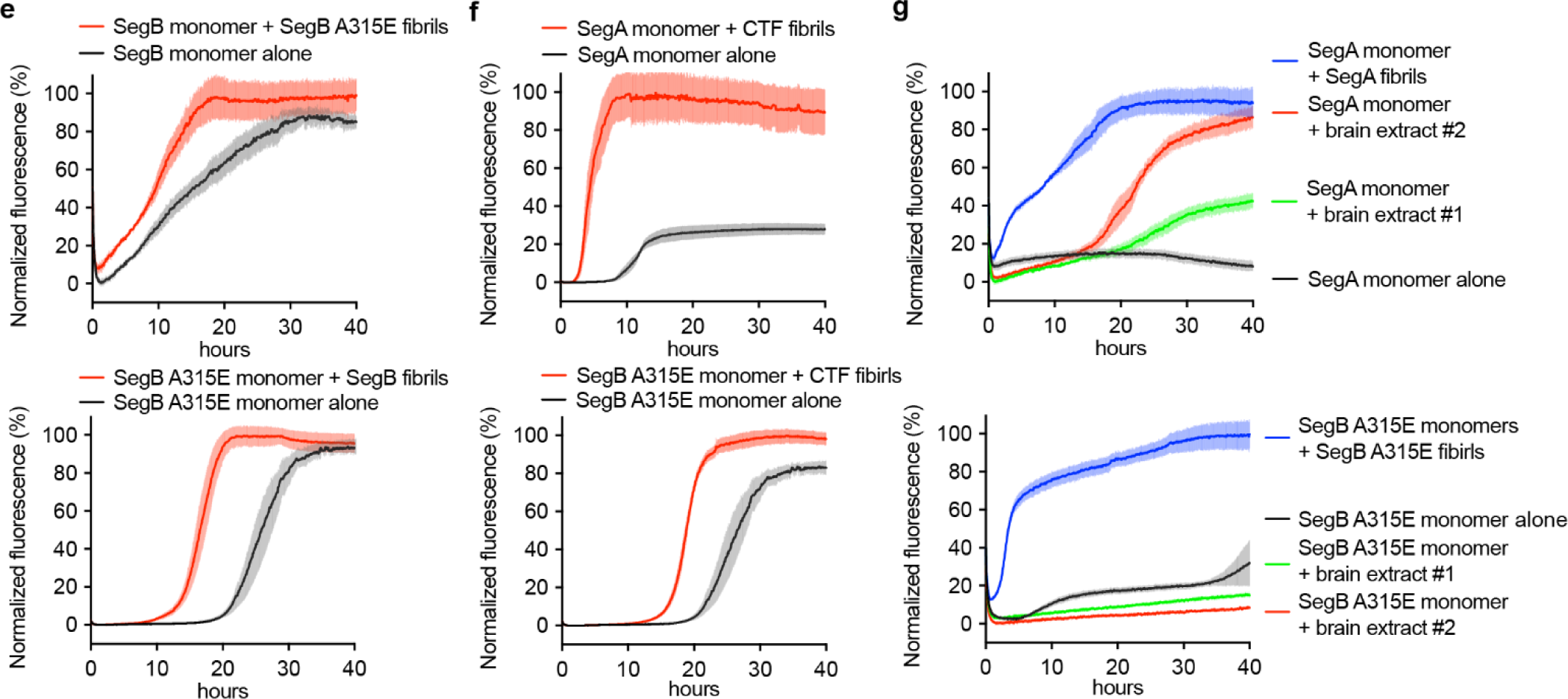
ThT seeding assays of TDP-43 segments. ThT aggregation curves of TDP-43 segments with or without seeding. Data are shown as mean ± s.d., n=4 independent experiments.

**Supplementary Figure 2.**
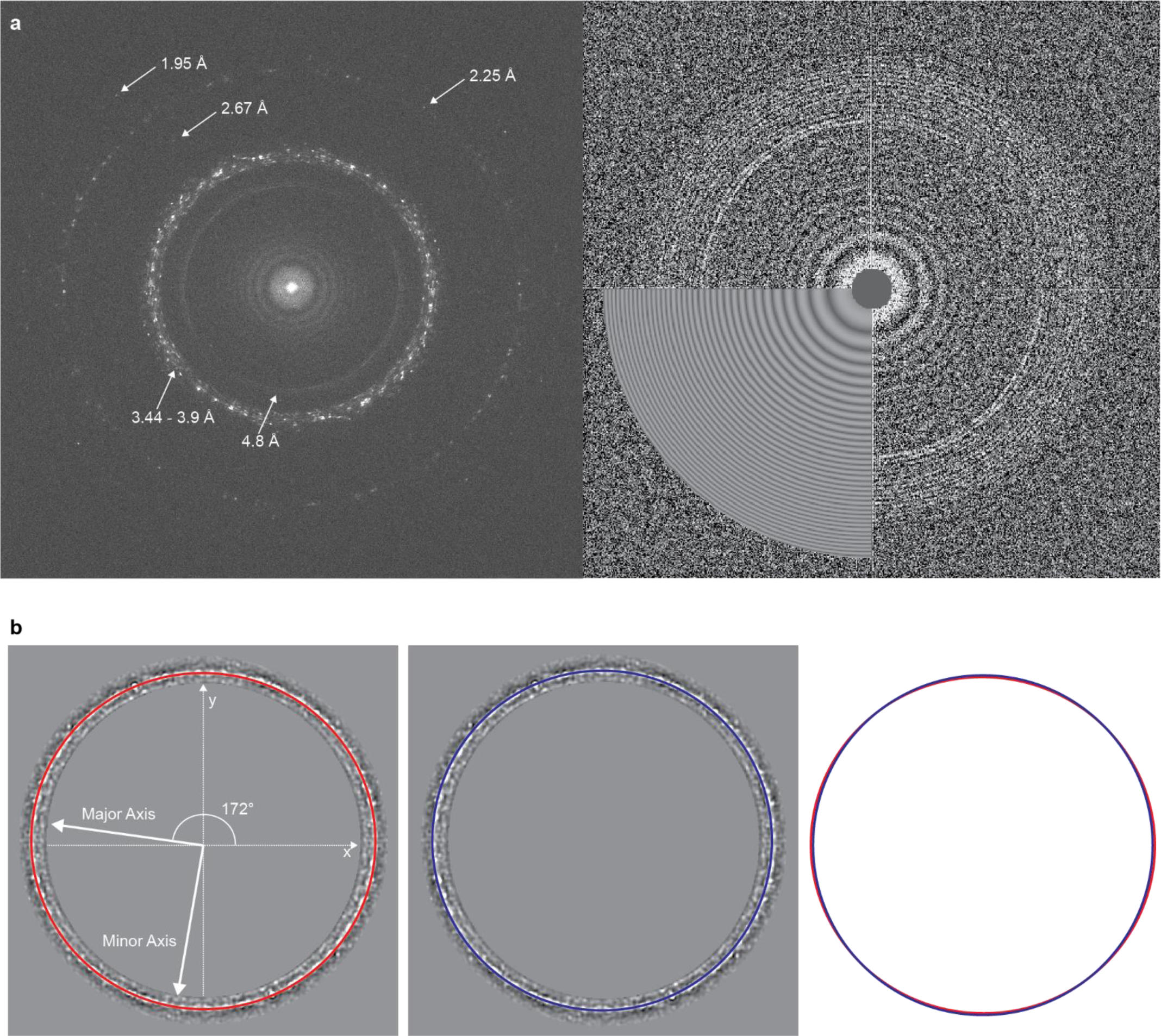
Alignment of Titan Krios microscope. **a**, (Left) Computed diffraction pattern from a micrograph containing crystalline ice with reflections visible to 1.95 Å demonstrating proper alignment of the microscope. (Right) Representative CTFFIND4 diagnostic image. **b**, Representative estimation and correction of anisotropic magnification distortion.

**Supplementary Figure 3.**
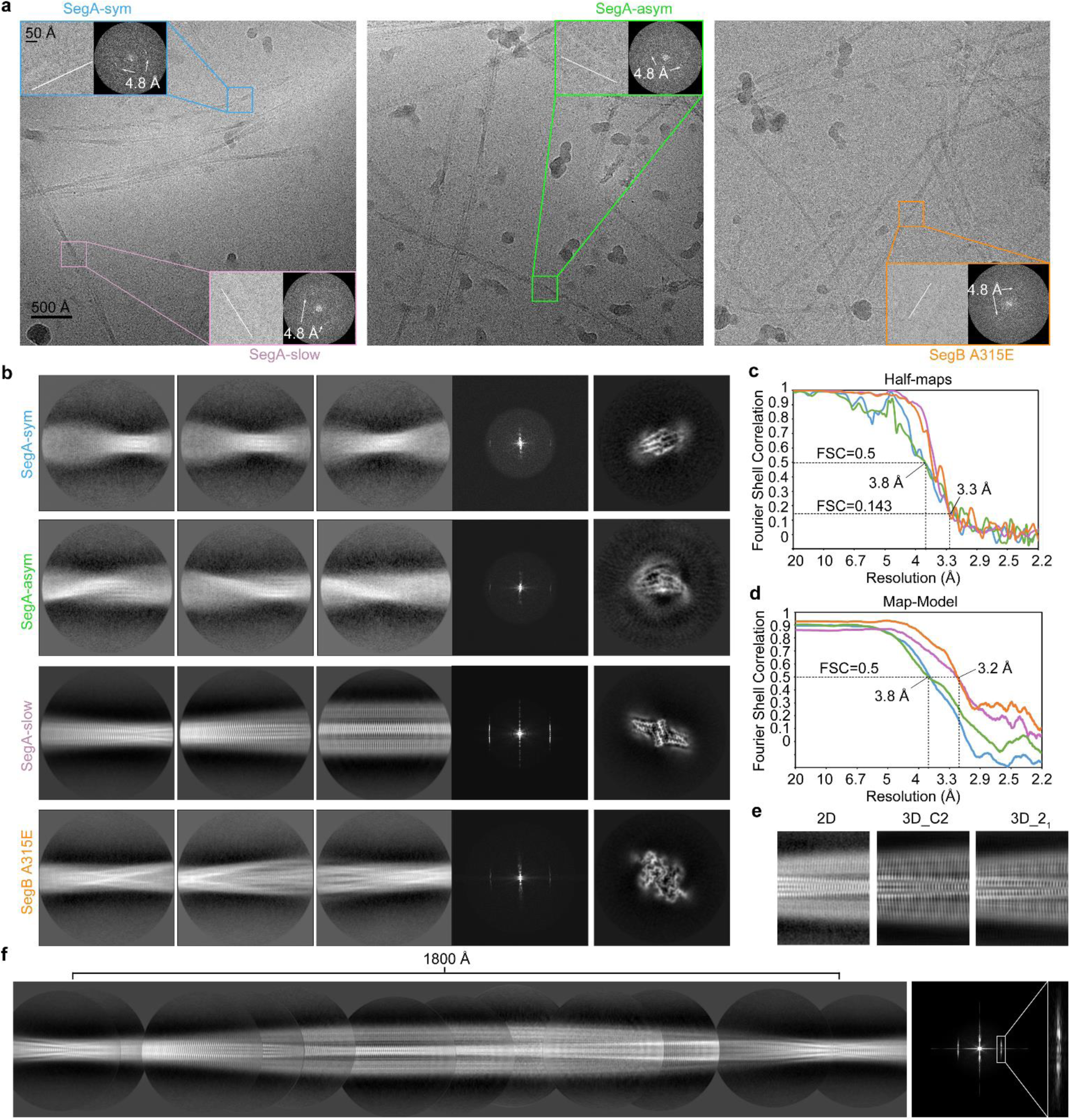
Cryo-EM data processing. **a**, Two (left and middle panel) and one (right) representative micrographs from data collection of SegA and SegB A315E, respectively. For each structure, a representative particle and its computed diffraction pattern are shown in the inset. The white line in each particle image indicates the direction of the fibril axis. The 4.8 Å reflection in the diffraction pattern corresponds to the helical rise. **b**, (left) Representative 2D class averages, (middle) computed diffraction pattern from 2D class, and (right) central slice from the final reconstruction of each structure. **c**-**d**, FSC curves between two half-maps (**c**) and the cryo-EM reconstruction and refined atomic model (**d**) for each structure, using the same color coding as in (a) and (b). **e**, Comparison of 2D class average (left) and projection of 3D reconstruction from the same orientation of the slow twister, when using C2 (middle) or pseudo-2_1_ (right) symmetry for reconstruction. **f**, Manually assembled full pitch of slow twister from 2D class averages and its computed diffraction pattern with the 4.8 Å region enlarged confirms the presence of C_n_ rotational symmetry due to a reflection at n=0.

**Supplementary Figure 4.**
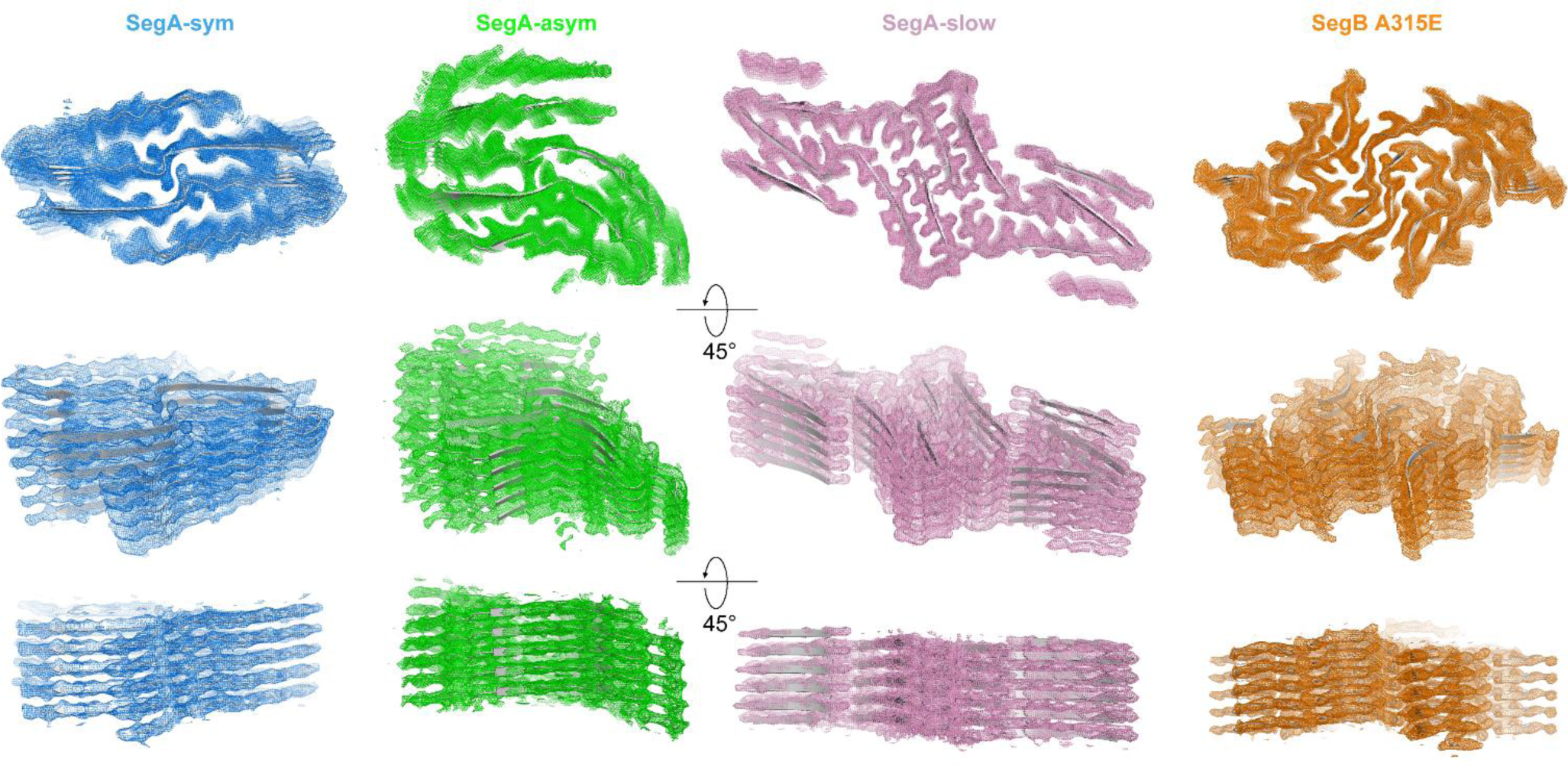
Density maps of TDP-43 fibril structures. Different views of the four structures reported in this study, with 5 layers of each structure shown. Density maps are shown in colored mesh and atomic model are shown in grey cartoons. Top-views demonstrate clear separation of beta-strands while tilted views demonstrate clear separation of the beta-sheets in the z-direction.

**Supplementary Figure 5.**
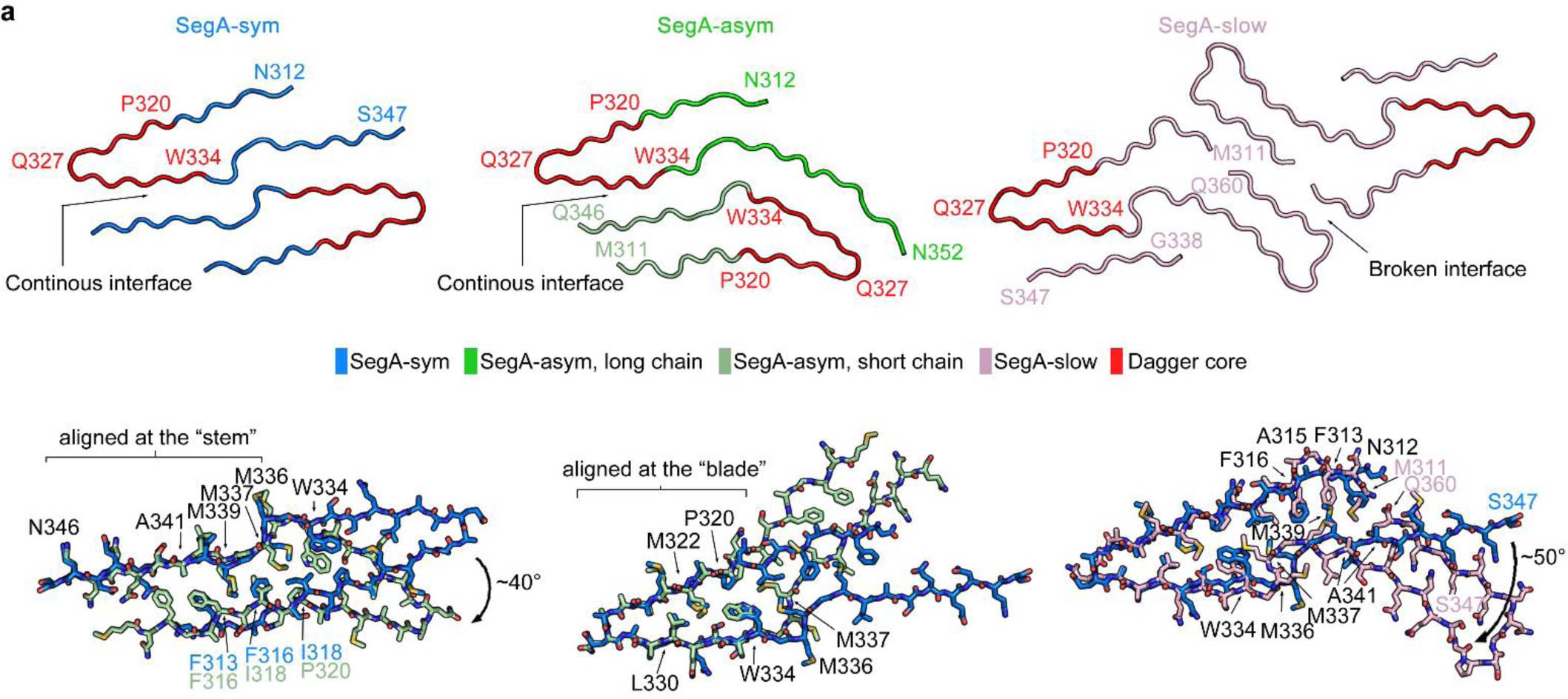
Detailed analysis in TDP-43 fibril structures. **a**, (Upper panels) Three polymorphs of SegA are shown in cartoon and colored according to the color key. The core region of the dagger-shaped fold (residue 320-334) is colored red. (Bottom panels) Pairwise superposition of the dagger-shaped fold. (Left and middle) SegA-sym vs. SegA-asym short chain aligned at stem and dagger, respectively; (right) SegA-sym vs. SegA-slow. (Detailed alignment parameters are listed in Supplementary Table 2).

**Supplementary Figure 5.**
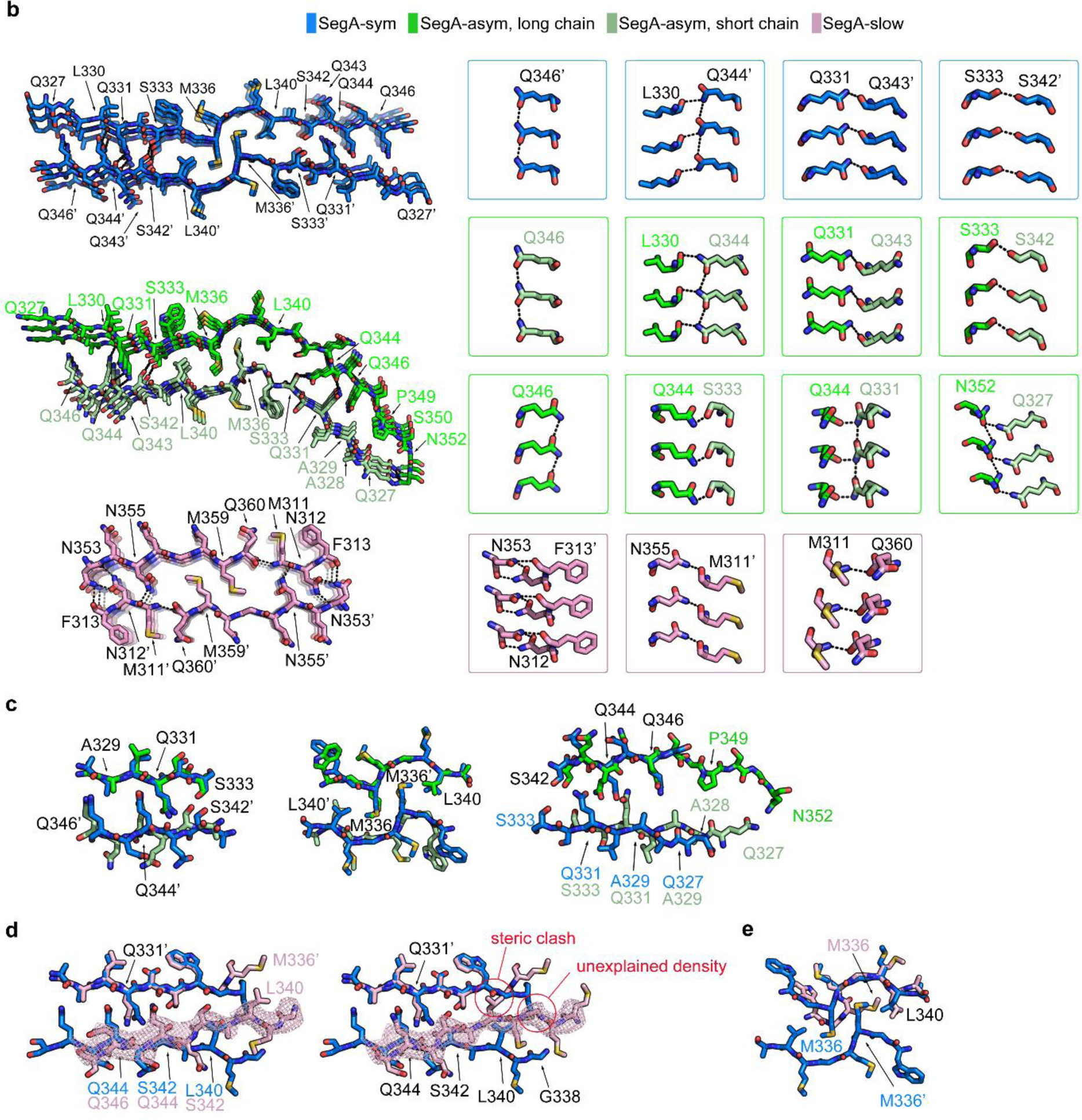
Detailed analysis in TDP-43 fibril structures. **b**, (Left column, from top to bottom) Cross-section of three layers of the continuous dimer interface in SegA-sym and SegA-asym, and broken interface in SegA-slow are shown. Side view of the detailed interactions of each interface are shown in the right columns. Hydrogen bonds are shown as black dashed lines with distance between 2.3-3.2 Å. **c**, Superposition of continuous interface in SegA-sym and SegA-asym. Because the 40° bend, we divided the continuous interface to three regions and perform superposition separately. **d**, Model of the SegA-slow continuous interface with 2-residue different (left) or same (right) registration as in SegA-sym. Notice that the clash at Leu340 and unexplained density at Gly338 makes same registration model unfavorable. **e**, Conformational change of Met336 in SegA-slow block the Met336-Leu340 pocket of continuous interface shown in SegA-sym. (Detailed alignment parameters are listed in Supplementary Table 2).

**Supplementary Figure 5.**
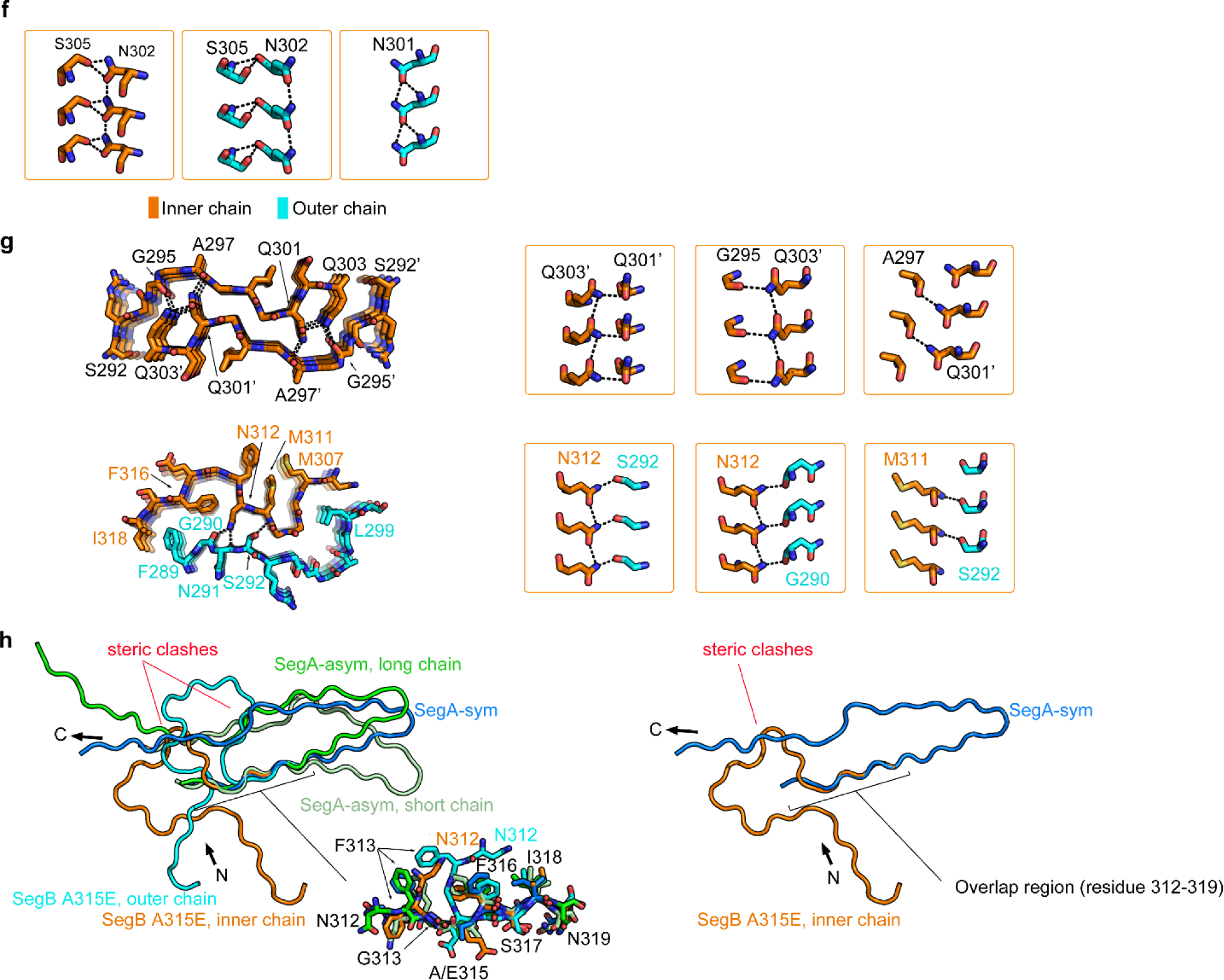
Detailed analysis in TDP-43 fibril structures. **f**, Side view of hydrophilic interactions in the head of the R-shaped fold in the SegB A315E. **g**, (Left column, from top to bottom) Cross-section of three layers of the homodimer and heterodimer interfaces in SegB A315E are shown. Side view of the detailed interactions of each interface are shown in the right columns. **h**, Superposition of SegA and SegB A315E main chain models shows TDP-43 cannot simultaneously form a dagger-shaped fibril and an R-shaped fibril because of steric clashes; it must form either one or other. The simplified comparison using only SegA-sym to represent the dagger-shaped fold and SegB A315E inner chain to represent the R-shaped fold are showing on the right panel. Color coding in this figure is the same as in Figure 2 and 3, hydrogen bonds are shown as black dashed lines with distance between 2.3-3.2 Å. Detailed alignment parameters are listed in Supplementary Table 2.

**Supplementary Figure 6.**
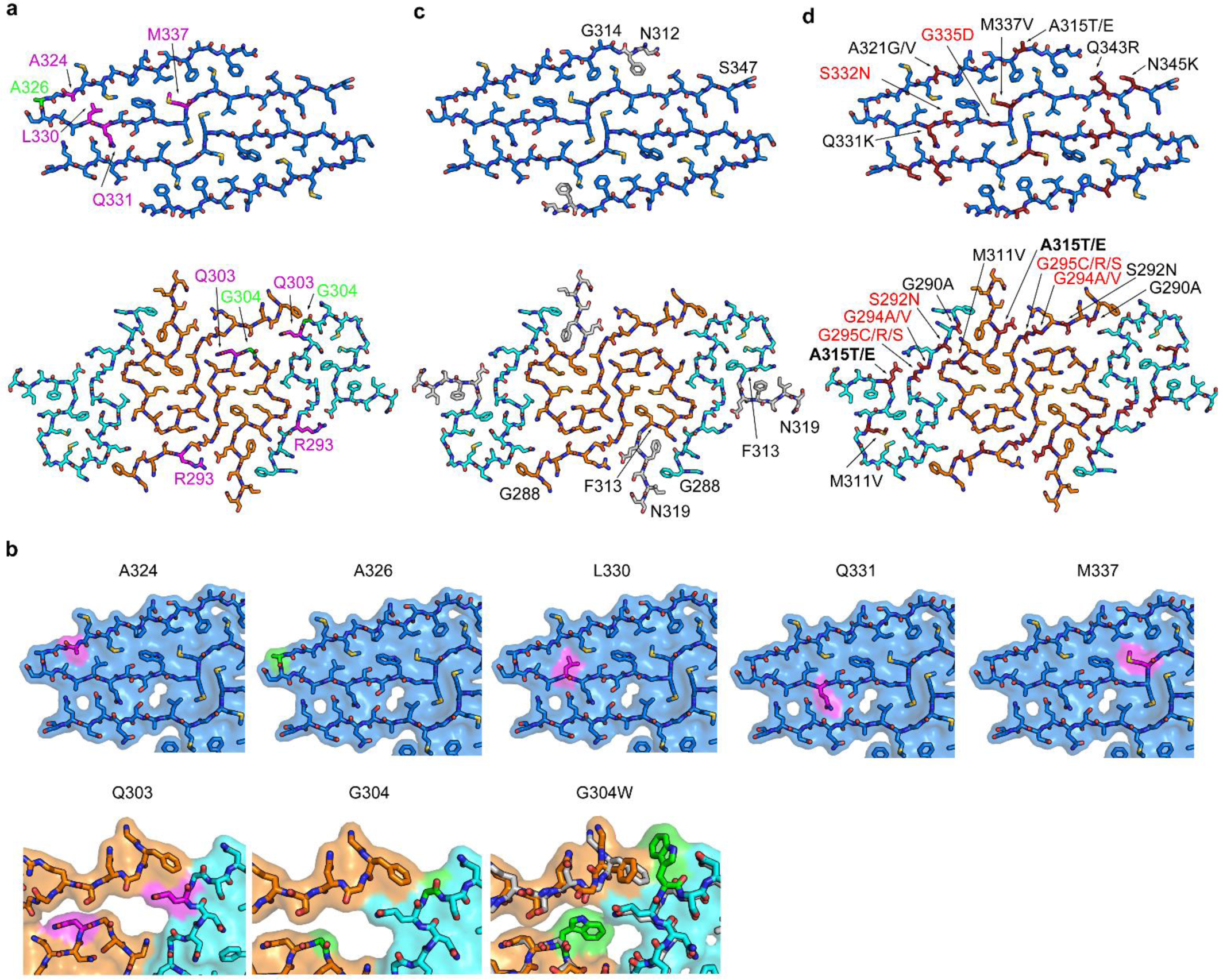
Mutation analysis of TDP-43 fibril structures. **a & b**, Residues tested in mutagenesis study in (top) SegA-sym and (bottom) SegB A315E. Residues that reduce pathological fragment TDP-CTF aggregation when mutated are shown in magenta and those that do not influence TDP-CTF aggregation when mutated are shown in green. Arg293 was mutated to glutamate and others were mutated to tryptophan. In (b), the models of both structures are shown in sticks and semitransparent surface mode. Notice that Ala324, Leu330, Gln331 and Met337 in SegA-sym and Gln303 in SegB A315E (also see Supplementary Figure 7) are in a tightly packed environment that does not tolerant a tryptophan mutation, consistent with our previous mutagenesis study where these mutations delayed the aggregation of the pathogenic TDP-43 fragment. In contrast, the Ala326 in SegA-sym and Gly304 in SegB A315E are in a relatively loose environment that is able to accommodate the tryptophan mutations, consistent with the result that A326W and G304W did not delay aggregation of the pathogenic TDP-43 fragment. For the G304W mutation, a plausible model of the R-shaped fold with G304W is shown in a separated panel. G304 from the inner chain and outer chain of SegB A315E were computationally mutated to tryptophan residues and phenix.real_space_refine was used to generate a model with acceptable rotamers and Ramachandran angles (Supplementary Table 3). The mutated model is shown in color and the non-mutated model is shown in gray. This model helps explain the tolerance of the G304W mutation in our previous study. **c**, Synthetic TDP-43 peptide 274-313 and 314-353 were reported to seed TDP-43 aggregation in cells, and the overlapping region of these two peptides in SegA-sym (left, 314-347) and SegB A315E (right, 288-313) structures were shown in color, whereas missing residues from the synthetic peptides are shown in grey. Notice that these two peptides preserve most of the dagger-shaped and the R-shaped fold making it likely that they adopt the same structures we report here. **d**, Positions of known hereditary mutations. The mutations compatible with the dagger-or R-shaped fold are colored black; the mutations potentially disruptive are colored red; the mutations favorable are bolded black.

**Supplementary Figure 7.**
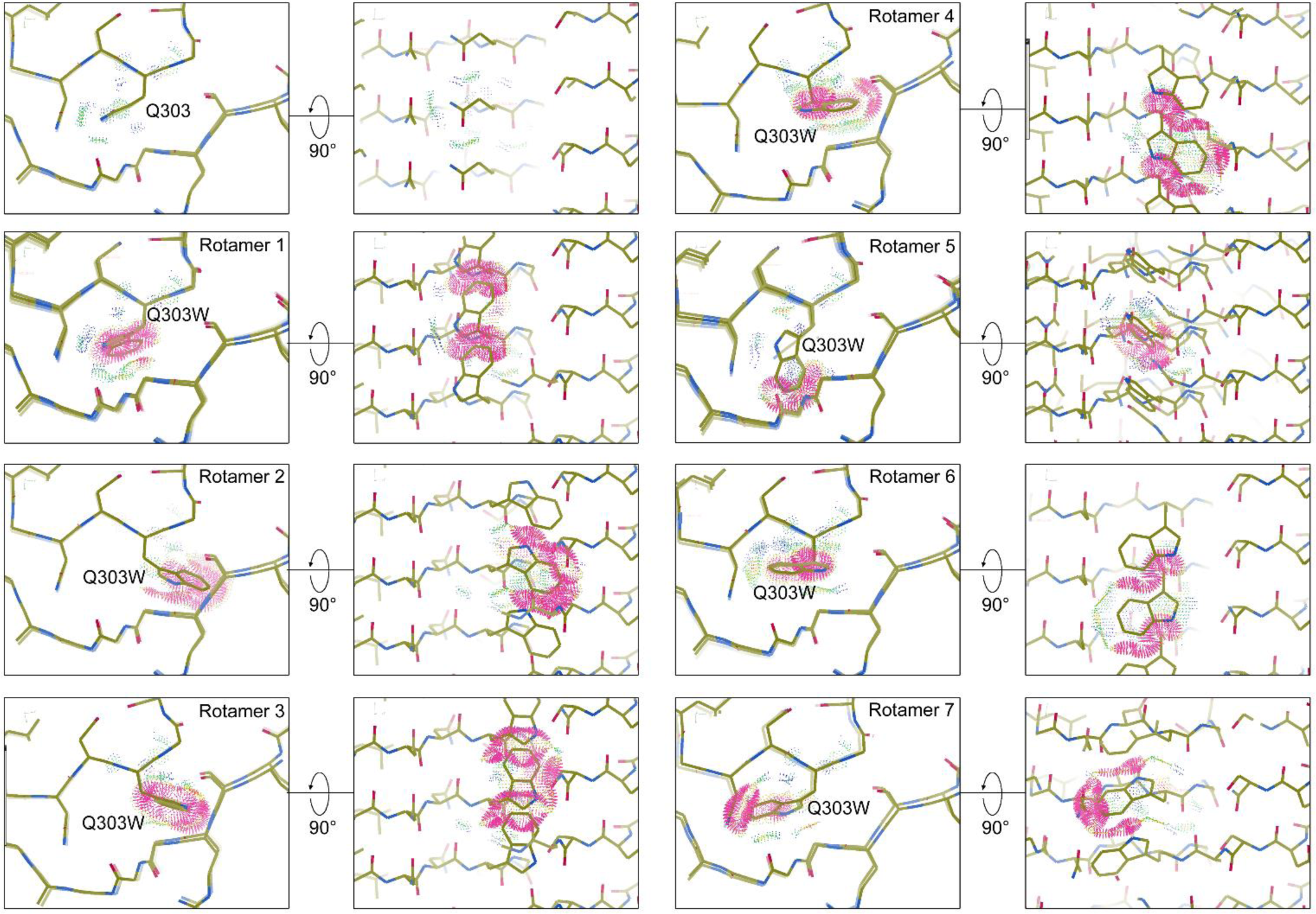
Space limitation of Q303W in the R-shaped fold. Gln303 in three continuous layers of the inner chain of the R-shaped fold was computationally mutated to tryptophan with all possible rotamers adopted. Steric clash was probed with COOT and displayed as red dots. Notice that the un-mutated Gln303 show no steric clash whereas all rotamers of Try303 show steric clash, indicating that Q303W is not compatible with the R-shaped fold.

**Supplementary Figure 8.**
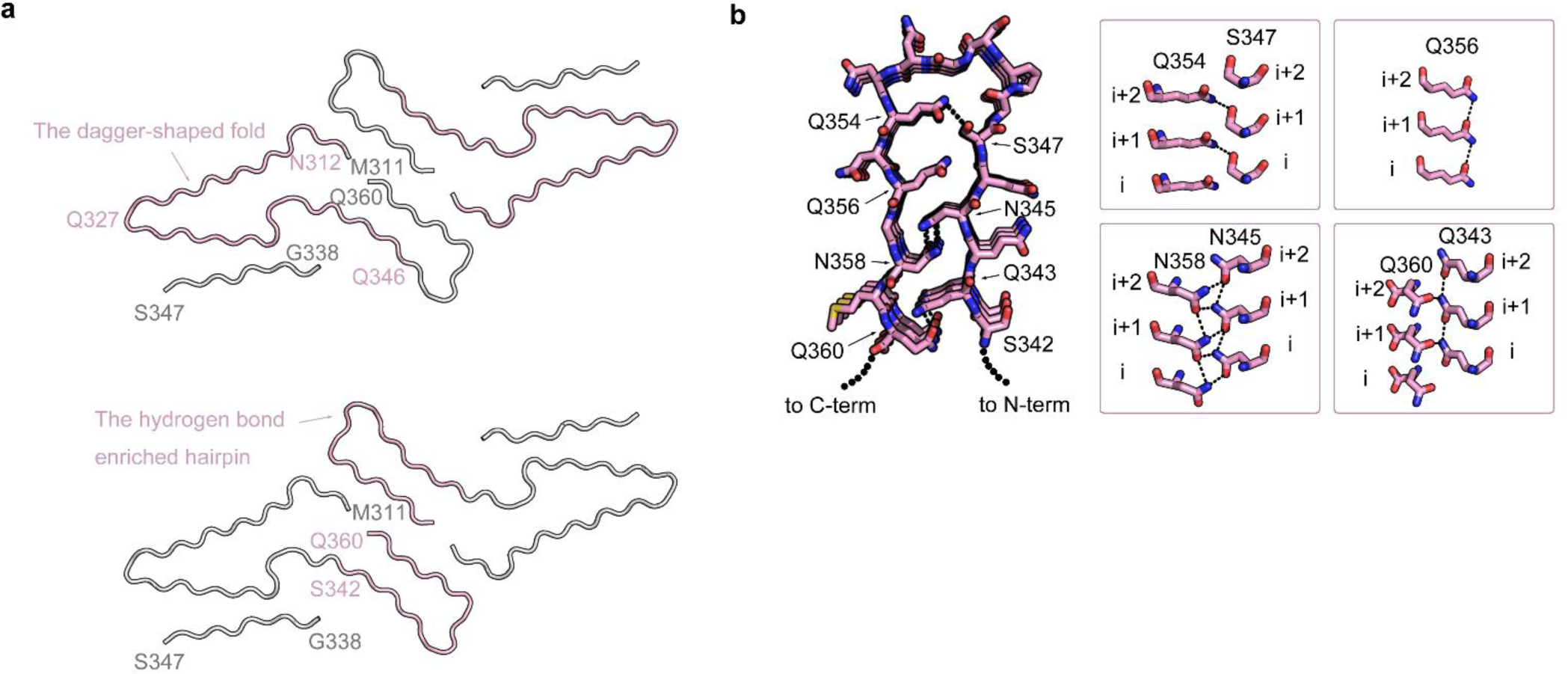
The dagger-shaped fold and the hydrogen bond enriched hairpin structure in SegA-slow. **a**, SegA-slow is shown in cartoon. The dagger-shaped fold (residue 312-346, upper panel) and the hydrogen bond enriched hairpin (residue 342-360, bottom panel) is colored pink, and the rest of the residues are colored gray. Note that both the dagger-shaped fold and the hydrogen bond enriched hairpin are individually compatible with full-length TDP-43 with free N-and C-terminals. **b**, Top view of hydrogen bond enriched hairpin structure in SegA-slow (left) and (right) side views of residues involved in hydrogen bond network.

## Supplemental Tables

**Supplementary Table 1.**
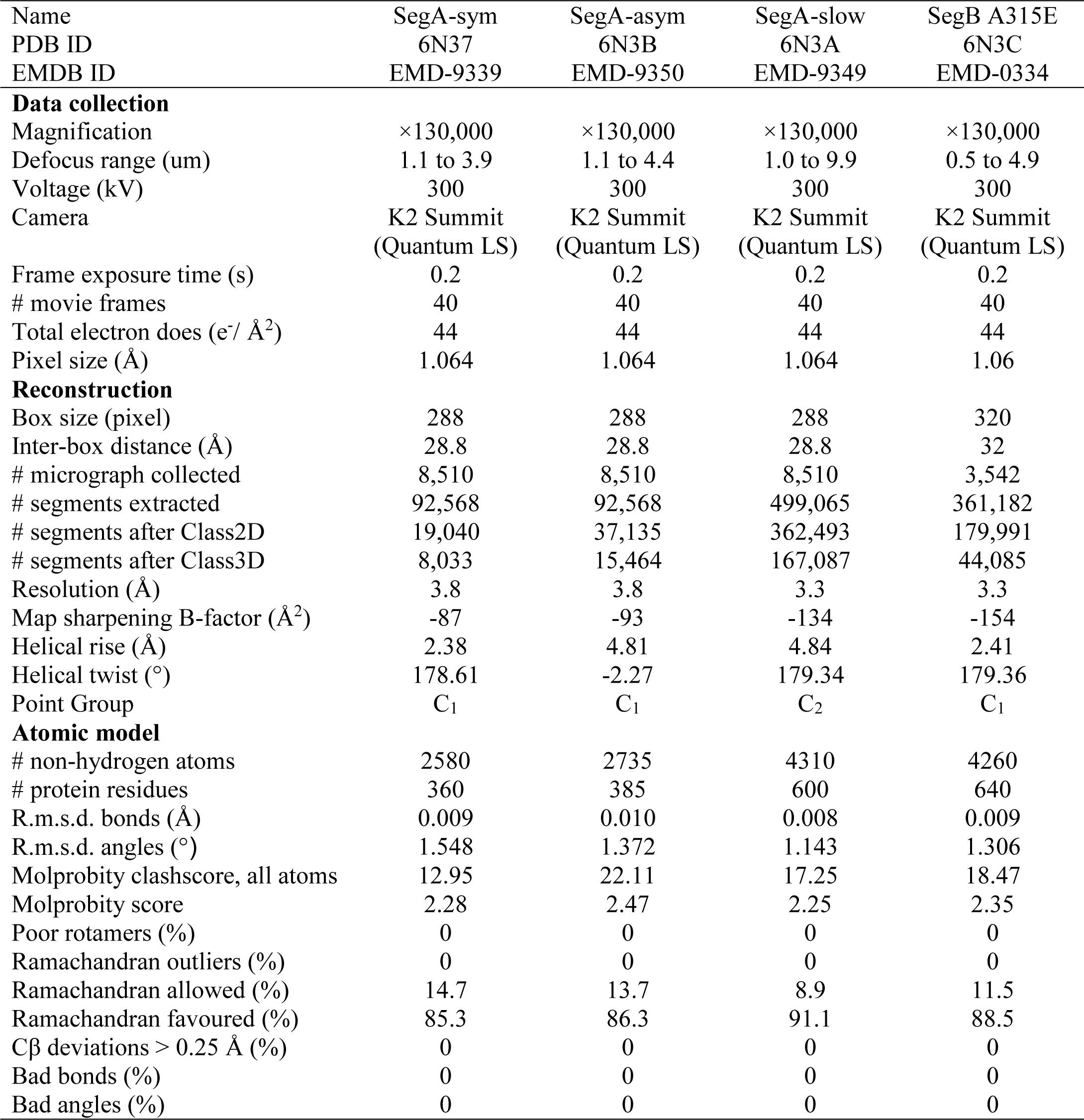
Cryo-EM data collection, refinement, and validation statistics.

**Supplementary Table 2.**
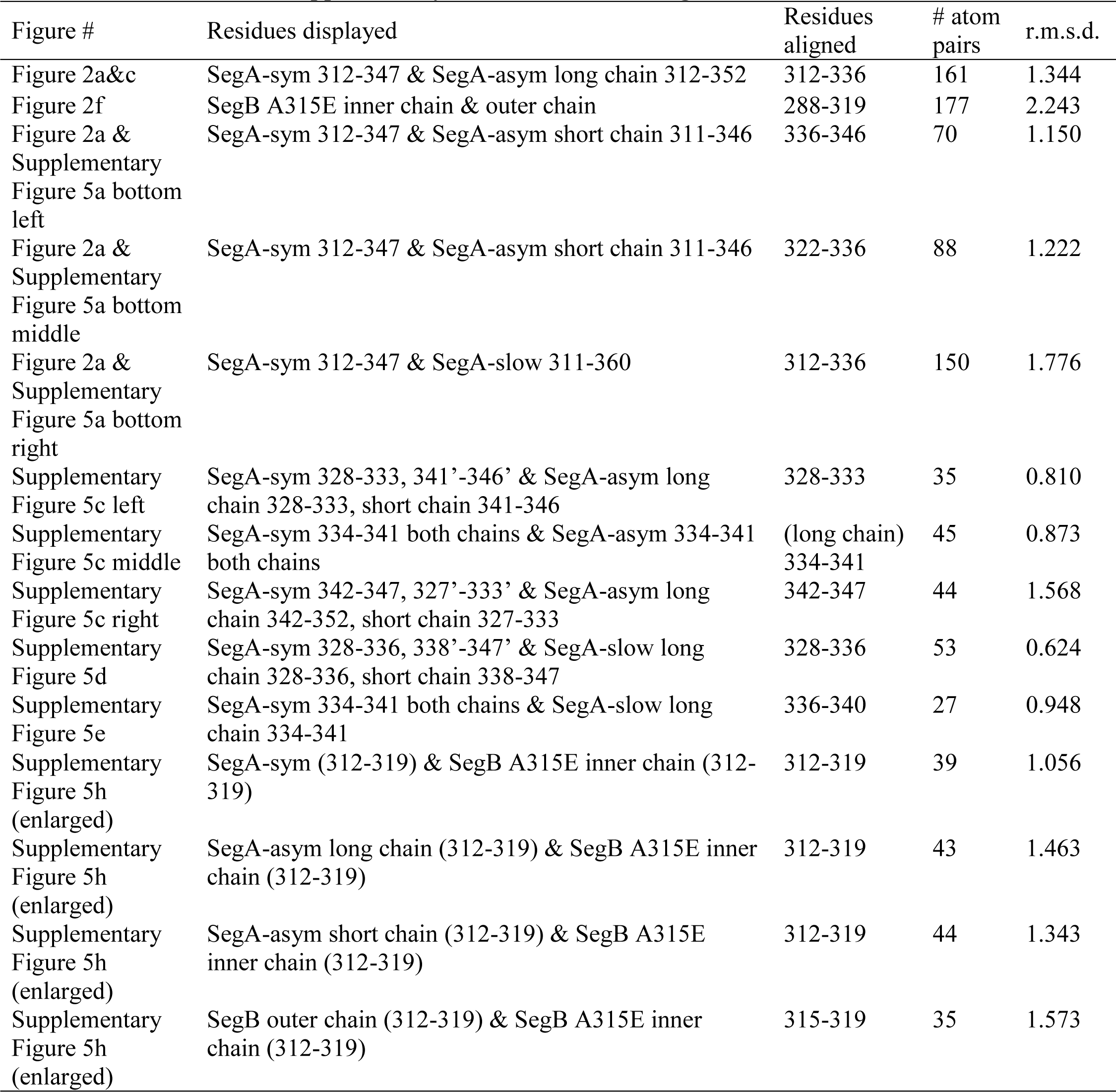
Structural alignment details.

**Supplementary Table 3.**
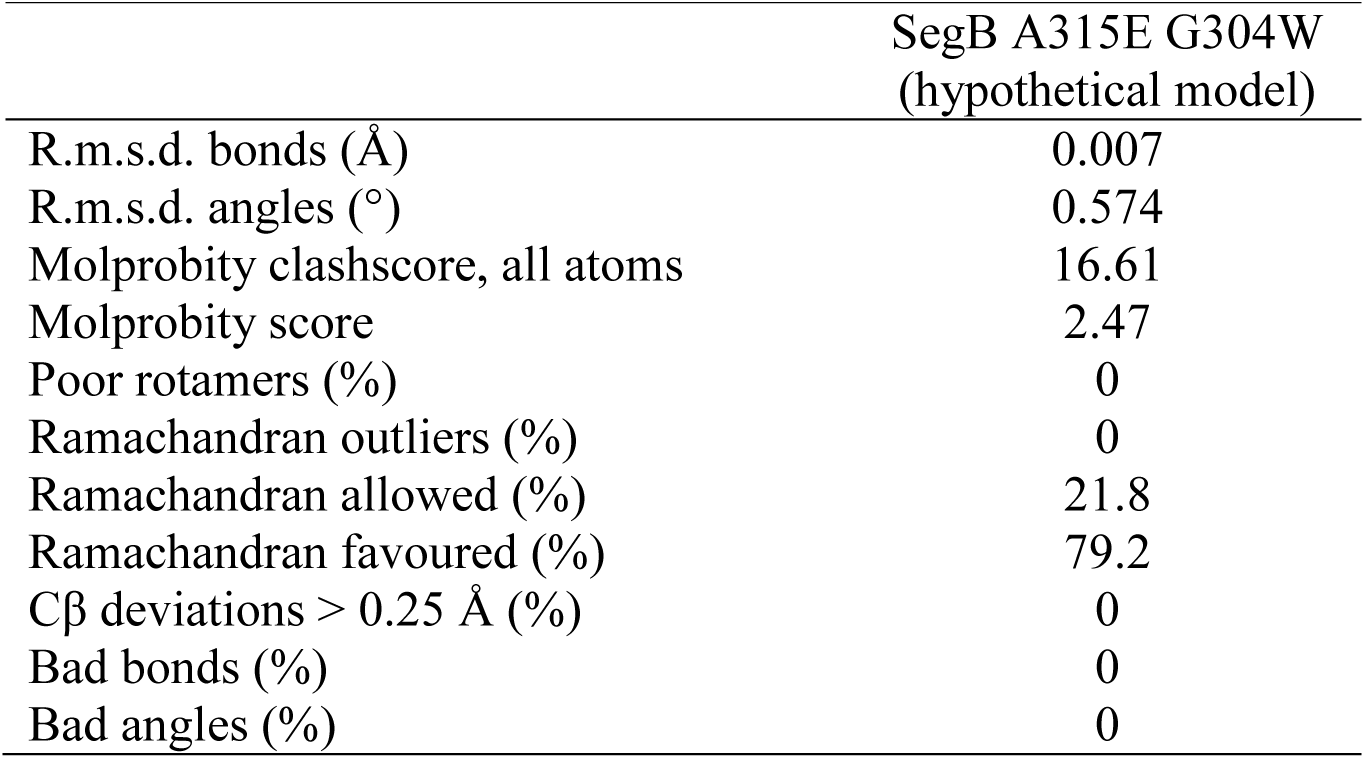
Validation statistics of the hypothetical SegB A315E G304W model.

|  | SegB A315E G304W<br>(hypothetical model) |
| --- | --- |
| R.m.s.d. bonds (Å) | 0.007 |
| R.m.s.d. angles (°) | 0.574 |
| Molprobity clashscore, all atoms | 16.61 |
| Molprobity score | 2.47 |
| Poor rotamers (%) | 0 |
| Ramachandran outliers (%) | 0 |
| Ramachandran allowed (%) | 21.8 |
| Ramachandran favoured (%) | 79.2 |
| Cβ deviations > 0.25 Å (%) | 0 |
| Bad bonds (%) | 0 |
| Bad angles (%) | 0 |

